# PRR14 organizes H3K9me3-modifed heterochromatin at the nuclear lamina

**DOI:** 10.1101/2022.05.17.492278

**Authors:** Anna A. Kiseleva, Yu-Chia Cheng, Cheryl L. Smith, Richard A. Katz, Andrey Poleshko

## Abstract

The eukaryotic genome is organized in three dimensions within the nucleus. Transcriptionally active chromatin is spatially separated from silent heterochromatin, a large fraction of which is located at the nuclear periphery. However, the mechanisms by which chromatin is localized at the nuclear periphery remain poorly understood. Here we demonstrate that Proline Rich 14 (PRR14) protein specifically organizes H3K9me3-modified heterochromatin at the nuclear lamina. We show that PRR14 dynamically associates with both the nuclear lamina and heterochromatin, and is able to reorganize heterochromatin in the nucleus of interphase cells independent of mitosis. We demonstrate that PRR14 can bind all isoforms of heterochromatin protein 1 (HP1). We characterize two functional HP1-binding sites within PRR14 that contribute to its association with heterochromatin. Results of fluorescent recovery after photobleaching (FRAP) and super-resolution imaging indicate that PPR14 forms an anchoring surface for heterochromatin at the nuclear lamina where it interacts dynamically with HP1-associated, H3K9me3-modified chromatin. Our study reveals the mechanism through which PRR14 tethers heterochromatin to the nuclear lamina and we propose a model of dynamic heterochromatin organization at the nuclear periphery.

## INTRODUCTION

The eukaryotic genome is organized at multiple scales in the nucleus. The genome is spatially arranged in subnuclear compartments so that transcriptionally active euchromatin is spatially segregated from silent heterochromatin (*1-4*). One example of spatially organized chromatin is transcriptionally silent heterochromatin sequestered at the periphery of the nucleus (*5, 6*). This precise organization of heterochromatin at the nuclear periphery contributes to many cellular functions such as mechanical force response, cell migration, signaling, transparency to light, and transcription control (reviewed in (*1-4, 7, 8*)). Multiple studies have shown that chromatin localization in the nucleus is often correlated with its transcriptional state: more often active when located in the nuclear interior and more often silenced when sequestered to the nuclear periphery (*9-11*). During development and lineage commitment, chromatin in the nucleus undergoes notable spatial reorganization with lineage-specific genes moving towards the nuclear interior while genes that are no longer required move to the nuclear lamina and become transcriptionally silenced (*5*). It has been demonstrated that correct spatial organization of heterochromatin at the nuclear lamina is essential for cell differentiation and lineage maintenance (*12-14*), however we know surprisingly little about the cellular mechanisms (epigenetic modifications, protein complexes, signaling pathways etc.) that orchestrate three-dimensional chromatin positioning within the nucleus in mammalian cells.

To date, only a few proteins have been shown to tether chromatin to the nuclear periphery (*15*). Lamin B Receptor (LBR) has been identified as an inner nuclear membrane protein that binds heterochromatin through the adapter protein Heterochromatin Protein 1 (HP1) (*16, 17*). Later studies identified the *C. elegans* CEC-4 protein as a membrane-associated heterochromatin tether (*13*) and mammalian Proline Rich 14 protein (PRR14) as a nuclear lamina-binding protein that also binds heterochromatin through the HP1 adapter (*18*). While mammalian LBR and PRR14 proteins bind heterochromatin through HP1, CEC-4 itself encodes an HP1-like domain (*13*), thereby obviating the need for an HP1 adapter. It is notable that all of the known peripheral heterochromatin tethers utilize a similar mechanism for heterochromatin binding.

Mammalian cells utilize three isoforms of HP1− HP1α, HP1β and HP1γ − which vary in their function and localization in the nucleus (*19-22*). HP1α and β isoforms are largely found in association with heterochromatin (*23*), while HP1γ is found in both heterochromatin and euchromatin compartments throughout the nucleus (*24, 25*). All HP1 isoforms are known to bind methylated Lys9 residue on Histone H3 tail (H3K9me) (*26, 27*). Both H3K9me2 and H3K9me3 histone modifications have been associated with heterochromatin, but specificity of interacting proteins has not been well defined.

LBR and CEC-4 are membrane-associated proteins that localize at the inner nuclear membrane and provide anchoring points for heterochromatin (*13, 28*). In contrast, PRR14 is a nuclear lamina-associated protein that is found primarily at the nuclear lamina with a much smaller fraction observed in the nucleoplasm. The observation that PRR14 localization is not strictly restricted to the nuclear periphery (*18, 29*) indicates that PRR14 association with the nuclear lamina is dynamic, and that the mechanism of chromatin tethering to the nuclear lamina via PRR14 could be different from that of LBR and CEC-4.

PRR14 was originally identified as an epigenetic repressor of gene expression (*30*). Further studies demonstrated that PRR14 localizes at the nuclear lamina, specifically in association with Lamin A/C (*18*). PRR14 has a 120-residue lamina-binding domain (LBD) that includes two independent lamina-binding motifs, both of which have been reported to be phosphoregulated (*29*). PRR14 also contains an HP1-binding motif in the N-terminal portion of the protein, and the PRR14-HP1 interaction has been shown to be required for PRR14 to bind chromatin (*31*). The C-terminus of the protein has a proposed regulatory function and includes a Tantalus domain with a Protein Phosphatase 2 Subunit Alpha (PP2A) binding motif. Interaction with PP2A is understood to regulate phosphorylation of the LBD and thus association of the protein with the nuclear lamina. While the domain structure of PRR14 has been described and several interacting proteins identified, the spatial dynamics of PRR14 in the nucleus and its ability to organize heterochromatin at the nuclear periphery has yet to be investigated.

Here we demonstrate that PRR14 functions at the nuclear lamina to organize heterochromatin that is, specifically, H3K9me3-modified, but not H3K9me2-modified. Our experimental results show that PRR14 associates dynamically with the nuclear lamina and heterochromatin, and is capable of reorganizing H3K9me3-modified heterochromatin in interphase, independent of mitosis. We assessed the ability of PRR14 to interact with the individual isoforms of heterochromatin protein 1 (HP1) and found an *in vivo* preference for HP1α and HP1β. In addition, we have identified a second HP1-binding motif in the N-terminus of the protein and show that it contributes to the association of PRR14 with heterochromatin, albeit to a lesser degree than the first PRR14 HP1-binding motif. Fluorescent recovery after photobleaching (FRAP) experiments demonstrated that PRR14 interactions with the nuclear lamina are relatively stable as compared with the more dynamic interactions of PRR14 with heterochromatin via HP1. Finally, our super-resolution imaging of PRR14 localization in the nucleus indicates that PRR14 is predominantly located at the nuclear lamina where, we predict, it provides an anchoring surface for HP1-associated, H3K9me3-modified chromatin. Combined, our results demonstrate the molecular mechanism of heterochromatin organization by PRR14 and suggest a dynamic model of heterochromatin tethering to the nuclear periphery.

## RESULTS

### PRR14 specifically tethers H3K9me3-modified heterochromatin to the nuclear lamina

*PRR14* encodes a protein with modular domains that bind to heterochromatin, via HP1, or to the nuclear lamina (Fig. 1A). Given this domain structure, it has been postulated that PRR14 functions to tether heterochromatin to the nuclear lamina (Fig. 1B) and this model is supported by loss-of-function experiments (*18*). It has not yet been determined whether PRR14 shows specificity for heterochromatin with particular histone modifications to be bound and positioned at the nuclear lamina. We tested the ability of PRR14 to tether H3K9me2- and H3K9me3-modified heterochromatin to the nuclear lamina in light of recent reports showing separate functions for H3K9me2 and H3K9me3 chromatin modifications (*12, 32, 33*).

**Figure 1.**
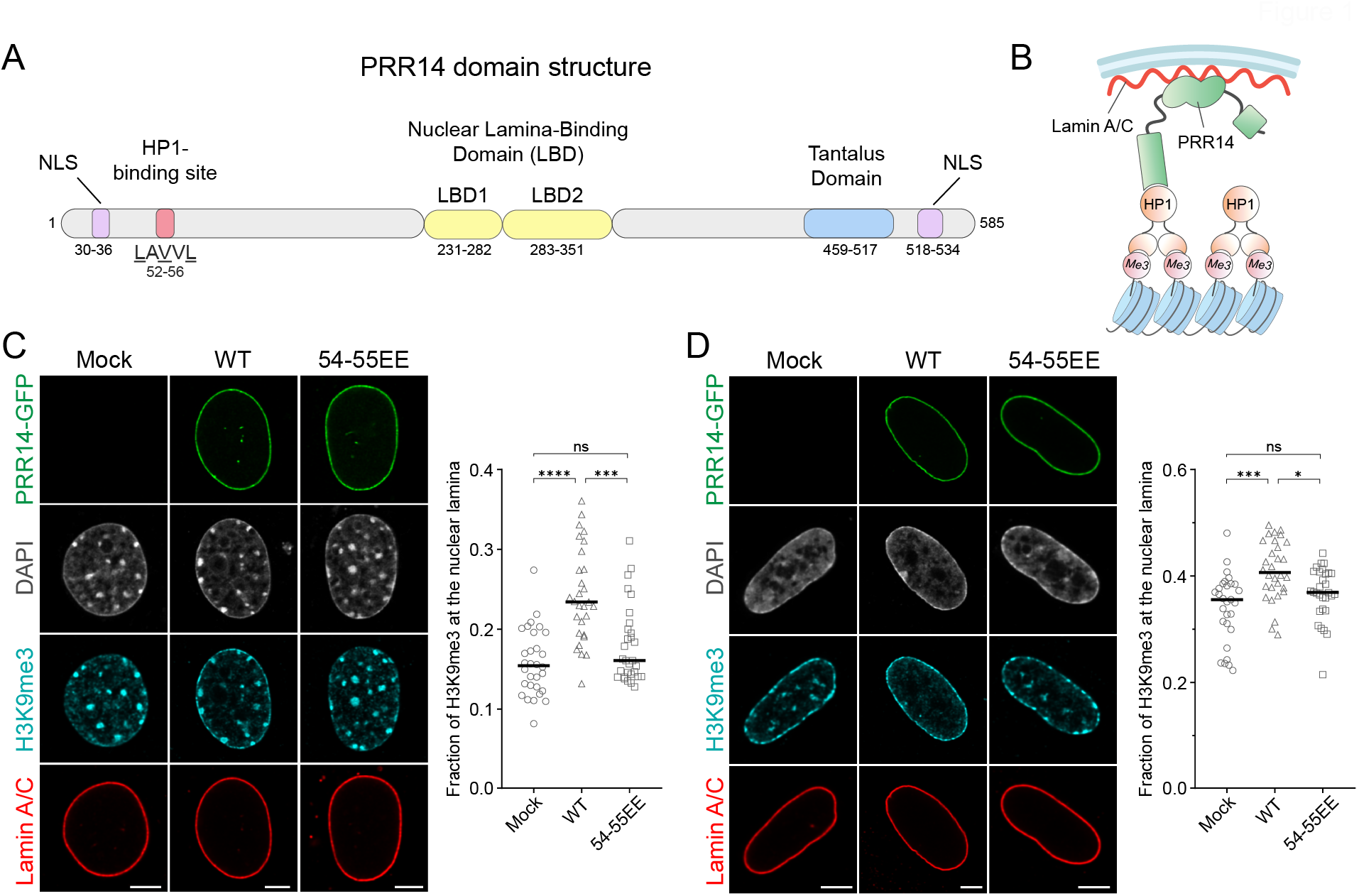
PRR14 specifically tethers H3K9me3-modified heterochromatin to the nuclear lamina. Schematic illustrations of **(A)** PRR14 domain organization, functional motifs and **(B)** mechanism of heterochromatin tethering to the nuclear lamina. Amino acid residues in (A) refer to human protein coordinates. NLS, nuclear localization signal. **(C-D)** Representative confocal images of (C) murine NIH/3T3 and (D) human IMR-90 cells expressing GFP-tagged PRR14 constructs (green): wildtype (WT) or mutant V54E, V55E (54-55EE) and stained for H3K9me3 (cyan) and Lamin A/C (red). DAPI counterstain shown in gray. Dot plots show the fraction of H3K9me3 signal at the nuclear lamina. n ≥ 30 cells per condition. Lines on dot plot show median values. Statistical analysis was performed using ANOVA Kruskal-Wallis test with Dunn’s multiple comparisons; ****p < 0.0001, ***p < 0.001, *p < 0.05, ns: not significant. Scale bars 5μm.

Exogenous expression of GFP-tagged PRR14 in two different cell types, murine NIH/3T3 and human IMR-90, altered the localization of H3K9me3-modified heterochromatin in the nucleus, significantly increasing that found at the nuclear lamina (Fig. 1C-D, Fig. S1A-B). This repositioning of H3K9me3-modified heterochromatin to the nuclear lamina was observed to be greater in NIH/3T3 cells, probably due to the lower baseline amount of H3K9me3 at the nuclear periphery prior to overexpression of PRR14. We did not observe any notable changes in the cellular amount of H3K9me3-modified heterochromatin following expression of PRR14-GFP constructs (Fig. S1C).

To determine if the relocalization of H3K9me3-modified heterochromatin depends on the HP1-binding domain of PRR14, we used a mutant form of PRR14 in which the HP1-binding motif (LAVVL) residues V54 and V55 were substituted to glutamic acid (54-55EE). In contrast to exogenous expression of wild-type PRR14, expression of the mutant PRR14 54-55EE construct did not alter the localization of H3K9me3-marked heterochromatin; rather cells expressing PRR14 54-55EE were similar to mock-transfected cells (Fig. 1C-D). Further, the ability of PRR14 to reposition heterochromatin was specific for that modified with H3K9me3. In cells exogenously expressing PRR14, we did not observe any repositioning effect for H3K9me2-marked heterochromatin (Fig. S2). Combined, these results demonstrate that PRR14 tethers H3K9me3-modified heterochromatin to the nuclear lamina through its HP1 binding domain.

### PRR14 organizes heterochromatin at the nuclear lamina in interphase nuclei independent of mitosis

PRR14 was previously reported to bind chromatin at early anaphase onset prior to nuclear lamina formation (*18*). Later in telophase, chromatin bound by PRR14 was observed to associate with newly formed nuclear lamina, and it was therefore proposed that PRR14 might play a role in re-establishing peripheral heterochromatin at mitotic exit (*31*). We sought to test whether mitosis is required for PRR14 to reorganize H3K9me3-modified heterochromatin in the nucleus.

NIH/3T3 cells expressing PRR14-GFP were assayed for changes in H3K9me3-modified heterochromatin localization following a double thymidine block to arrest the cell cycle and prevent mitosis (Fig. S3, see Methods). PRR14 overexpression in both cycling and arrested cells resulted in a significant increase in the fraction of H3K9me3-modified heterochromatin at the nuclear lamina with no significant difference between untreated and thymidine-treated cells (Fig. 2). Similar levels of H3K9me3 at the nuclear periphery of both arrested and cycling cells with PRR14 overexpression suggests that PRR14 can tether heterochromatin in interphase nuclei and organize heterochromatin at the nuclear lamina independent of mitosis.

**Figure 2.**
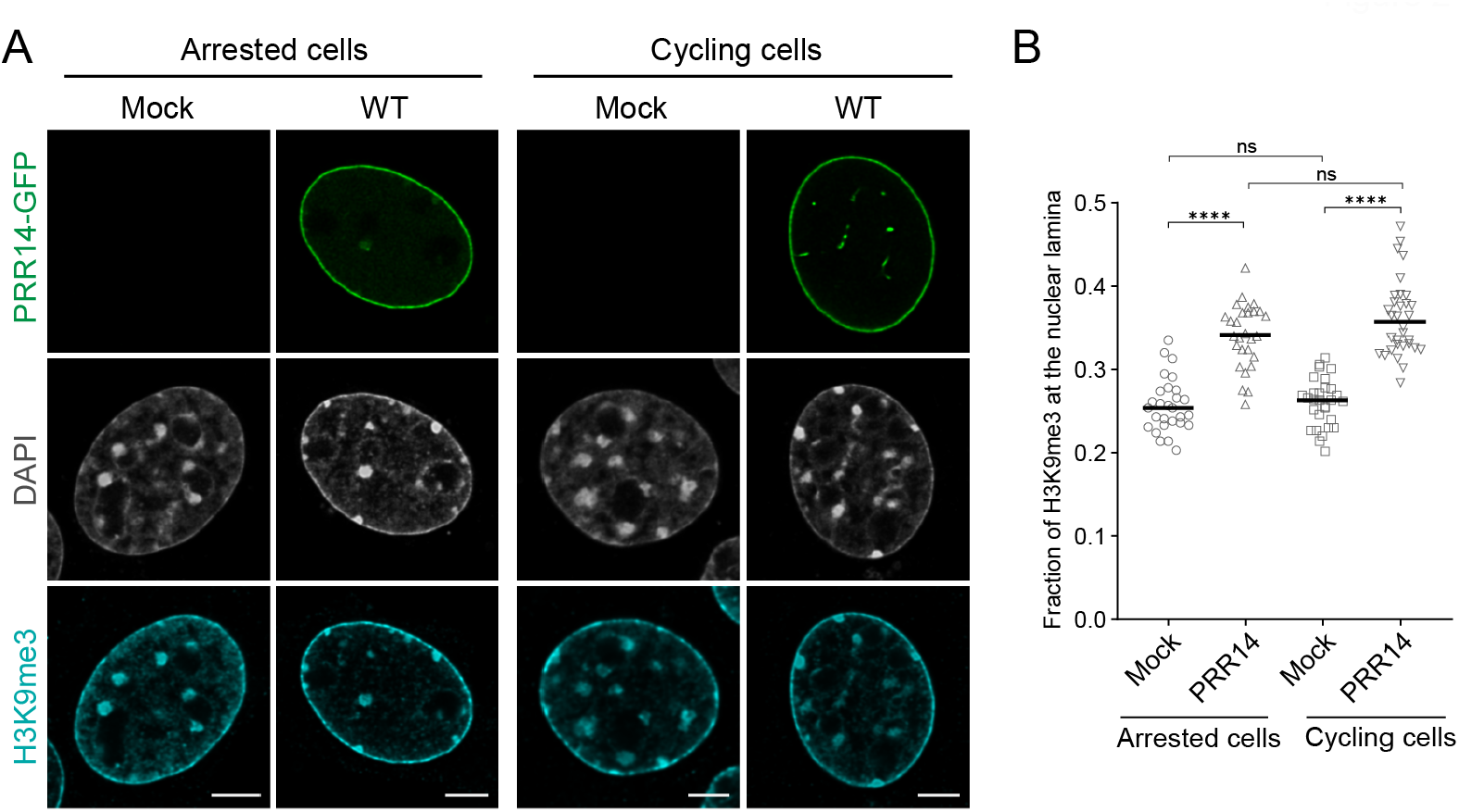
PRR14 organizes H3K9me3-modified heterochromatin at the nuclear lamina in interphase nuclei independent of mitosis. (A) Representative confocal images of cell cycle-arrested and cycling murine NIH/3T3 cells expressing WT GFP-tagged PRR14 constructs (green), stained for H3K9me3 (cyan). DAPI counterstain shown in gray. (B) Dot plot shows the fraction of H3K9me3 signal at the nuclear lamina. Lines on dot plot show median values. n ≥ 30 cells per condition. Statistical analysis was performed using ANOVA Kruskal-Wallis test with Dunn’s multiple comparisons; ****p < 0.0001, ns: not significant. Scale bars 5μm.

### PRR14 N terminus includes two conserved HP1-binding motifs

The LAVVL motif at the N-terminus of PRR14 (aa 52-56) is a variation of HP1-binding motif <u>L</u>x<u>V</u>x<u>L.</u> Substitution of key residues within this motif – V54E, V55E (LAEEL) – results in a dramatic decrease of PRR14 association with chromatin (Fig. 1) (*18*). Through motif scanning of the remainder of the protein, a second evolutionary conserved LxVxL sequence was identified within PRR14: LVVML, at amino acids 153-157 (Fig. 3A), just beyond the region that was originally implicated as the chromatin-binding domain (aa 1-135, (*18*)). We sought to determine if this site could also contribute to heterochromatin binding.

**Figure 3.**
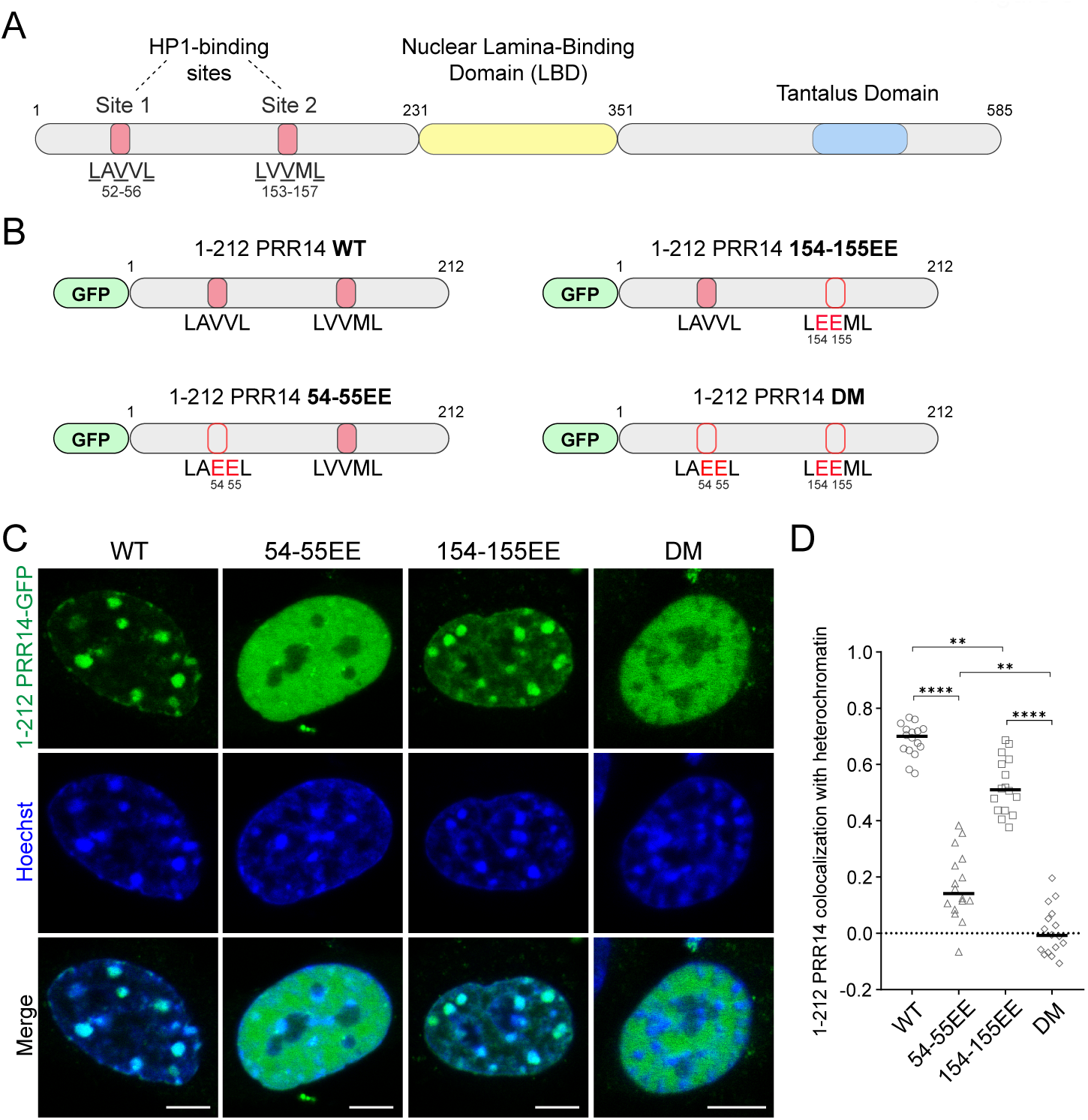
PRR14 interacts with heterochromatin primarily through its HP1-binding site 1. **(A)** Schematic illustrations of PRR14 domain organization and HP1-binding site motifs: LAVVL (aa 52-56) and LVVML (aa 153-157). **(B)** Schematic representation of GFP-tagged PRR14(1-212) fragments including the heterochromatin-binding domain of PRR14 with indicated amino acid substitutions in key residues of HP1-binding motifs. DM, double mutant. **(C)** Representative confocal images of murine NIH/3T3 cells expressing indicated WT or mutant GFP-tagged PRR14(1-212) constructs (green), counterstained with Hoechst (blue). **(D)** Dot plot shows the Pearson’s correlation of Hoechst staining and GFP-PRR14 signal for each PRR14(1-212) construct indicating degree of colocalization of PRR14 with heterochromatin regions. n ≥ 16 cells per condition. Lines on the dot plot show median values. Statistical analysis was performed using ANOVA Brown-Forsythe and Welch test with Dunnett’s multiple comparisons; ****p < 0.0001, **p < 0.01. Scale bars 5μm.

To compare the functions of the first and putative second HP1-binding motifs of PRR14, we made GFP-tagged PRR14 constructs encoding only the N-terminal 212 amino acids of the protein that terminate before the lamina-binding domain (LBD) (Fig. 3B). In addition to a wild-type version, we made GFP-tagged PRR14 1-212 constructs coding for amino acid substitutions in the first (V54E, V55E), the second (V154E, V155E), or both HP1-binding motifs (V54E, V55E, V154E, V155E), denoted 54-55EE; 154-155EE; and double mutant (DM), respectively (Fig. 3B).

We introduced the wild-type and mutant 1-212 fragments of PRR14 into NIH/3T3 cells, and as expected, the wild-type fragment showed consistent colocalization in the nucleus with chromocenters and other heterochromatic regions known to be marked with the H3K9me3 histone modification (*34*) (Fig. 3C). Further, the PRR14 1-212 fragment, which lacks the Lamina-Binding Domain (LBD) was, as predicted, not observed at the nuclear periphery (Fig. 3C).

Compared with WT PRR14 1-212 fragment, expression of the HP1 binding site 1 mutant version of PRR14 1-212 (54-55EE) resulted in a dramatic decrease, but not complete loss of PRR14 co-localization with heterochromatin as visualized by Hoechst staining. We observed a diffuse localization of this 54-55EE mutant in the nucleus and minimal colocalization with heterochromatin (Fig. 3C-D). In contrast, the HP-binding site 2 mutant (154-155EE) retained some association with heterochromatin, but it was significantly reduced compared with WT (Fig. 3C-D). PRR14 1-212 with mutations of both HP1 binding sites (DM) showed a complete absence of colocalization with heterochromatin (Fig. 3C-D). Taken together, these results demonstrate that HP1-binding site 1 plays the major role in heterochromatin binding, while HP1-binding site 2 contributes to a lesser degree to PRR14 interactions with heterochromatin. This is consistent with the observations that both PRR14 54-55EE and PRR14 DM lack the ability to efficiently tether heterochromatin to the nuclear lamina (Fig. 1C-D, Fig. S4).

To further examine the influence of PRR14 HP1 binding sites on heterochromatin interactions, we performed fluorescent recovery after photobleaching (FRAP) experiments. We probed the association of the PRR14 1-212 fragment with the large, H3K9me3-marked heterochromatin regions found at chromocenters in murine cells (*34*). There was a rapid recovery of the GFP-PRR14 signal following photobleaching of a single chromocenter in cells expressing the wild-type PRR14 1-212 fragment (Fig. 4A, Fig. S5). Normalized fluorescent recovery at photobleached regions was very high, with only about 15% average immobile fraction of WT PRR14 1-212 fragment on heterochromatin (Fig. 4B-D). The average recovery half-time for WT PRR14 1-212 fragment was 4.5 seconds (Fig. 4C) indicating a very rapid exchange and dynamic association of PRR14 with heterochromatin.

**Figure 4.**
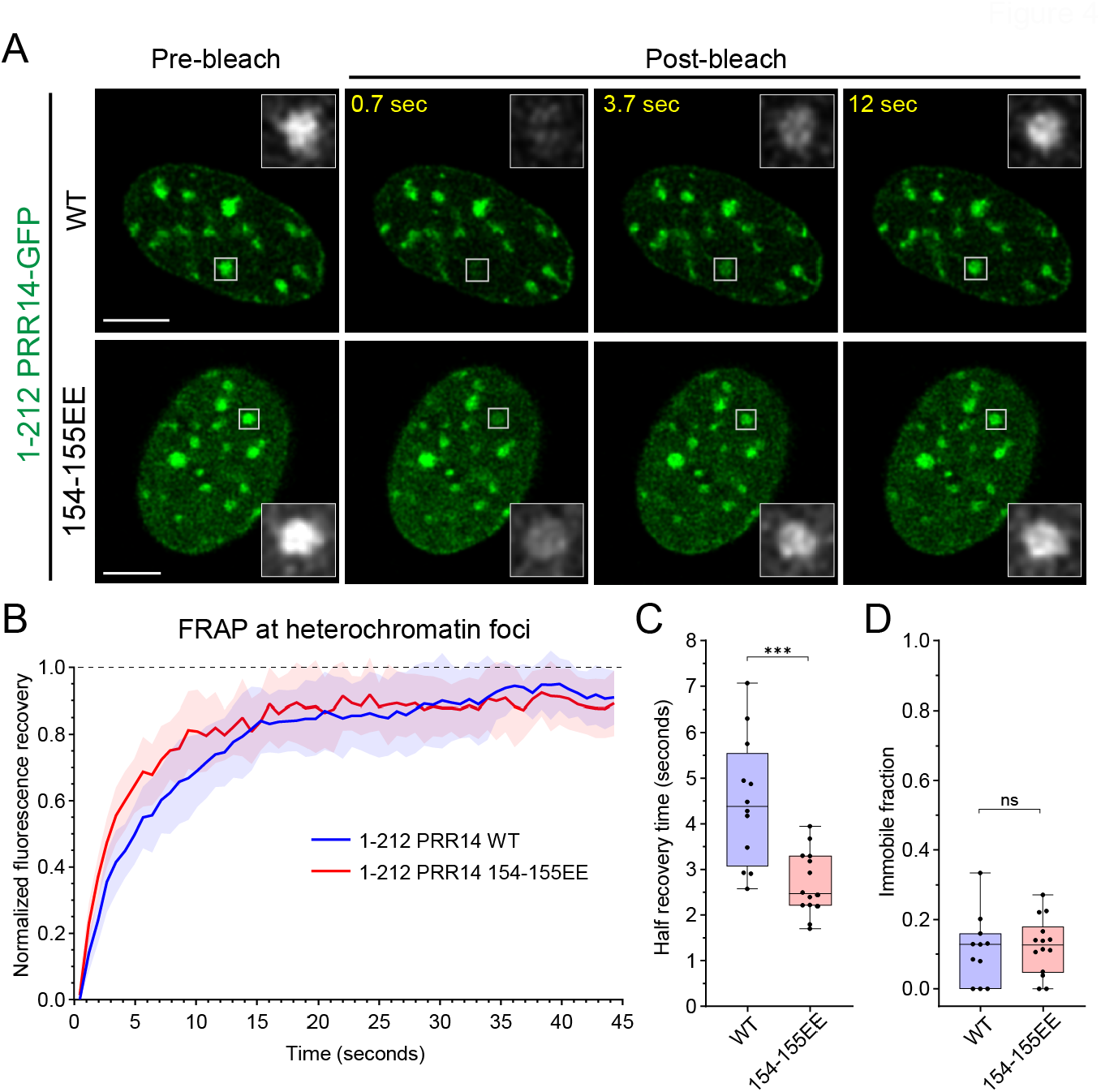
PRR14 HP1-binding site 2 stabilizes PRR14-heterochromatin interactions. **(A)** Representative confocal images of fluorescent recovery after photobleaching (FRAP) assay for PRR14(1-212) WT and 154-155EE mutant constructs. White boxes indicate the bleached area which are shown as magnified, grayscale images. **(B)** Line graph shows normalized fluorescent recovery over time after photobleaching in the areas indicated in (A) in cells expressing PRR14(1-212) WT (blue) or 154-155EE mutant (red). Line graph shows mean values with standard deviations displayed as shading. **(C)** Box plots show distributions of recovery half-times for indicated PRR14 (1-212) constructs. **(D)** Box plots show distributions of immobile fractions for indicated constructs. n ≥ 12 cells per condition. Box plots show median, 25th and 75th percentiles. Whiskers show minimum to maximum range. Statistical analysis was performed using Mann-Whitney test. ***p < 0.001, ns: not significant. Scale bars 5μm.

While the 1-212 PRR14 constructs with HP1-binding site mutations (54-55EE and DM) show little to no enrichment at heterochromatin loci (Fig. 3), we were able to measure those of the HP1-binding site 2 mutant (154-155EE) and compare its dynamics with the WT PRR14 fragment. The 1-212 PRR14 154-155EE mutant displayed heterochromatin association similar to that of WT but with a significantly faster average recovery half-time of 2.5 seconds (Fig. 4, Fig. S5). Thus, the interaction of PRR14 154-155EE with heterochromatin is even more dynamic than that of WT PRR14, consistent with a role for the second HP1-binding site in stabilizing the overall PRR14-chromatin association. The observed dynamics of PRR14 interaction with heterochromatin through its two HP1-binding sites is similar to that reported for another HP1-binding protein, SENP7, which displays evidence of two HP1-binding sites shown to contribute to the association of SENP7 with heterochromatin (*35*).

### PRR14 binding preference for HP1 isoforms

Given PRR14 chromatin binding is mediated by its interactions with HP1 proteins, we next sought to determine whether PRR14 interacts preferentially with any of the HP1 isoforms. Since the exogenous expression of full-length PRR14 resulted in relocalization of H3K9me3-modified heterochromatin to the nuclear lamina, we asked whether overexpression of full-length PRR14 led to an increase in the amount of any of the HP1 isoforms at the nuclear periphery. In murine NIH/3T3 cells transfected with full-length PRR14, we examined the localization of each of the HP1 isoforms. We observed a significant increase of both HP1α and HP1β isoforms at the nuclear lamina in PRR14-overexpressing cells as compared to mock transfected cells (Fig. 5A, B, Fig. S6). HP1γ, which is less tightly associated with heterochromatin and known to organize euchromatin (*24, 25*), but overexpression of PRR14 resulted in an observable change in HP1γ localization (Fig. 5A, B).

**Figure 5.**
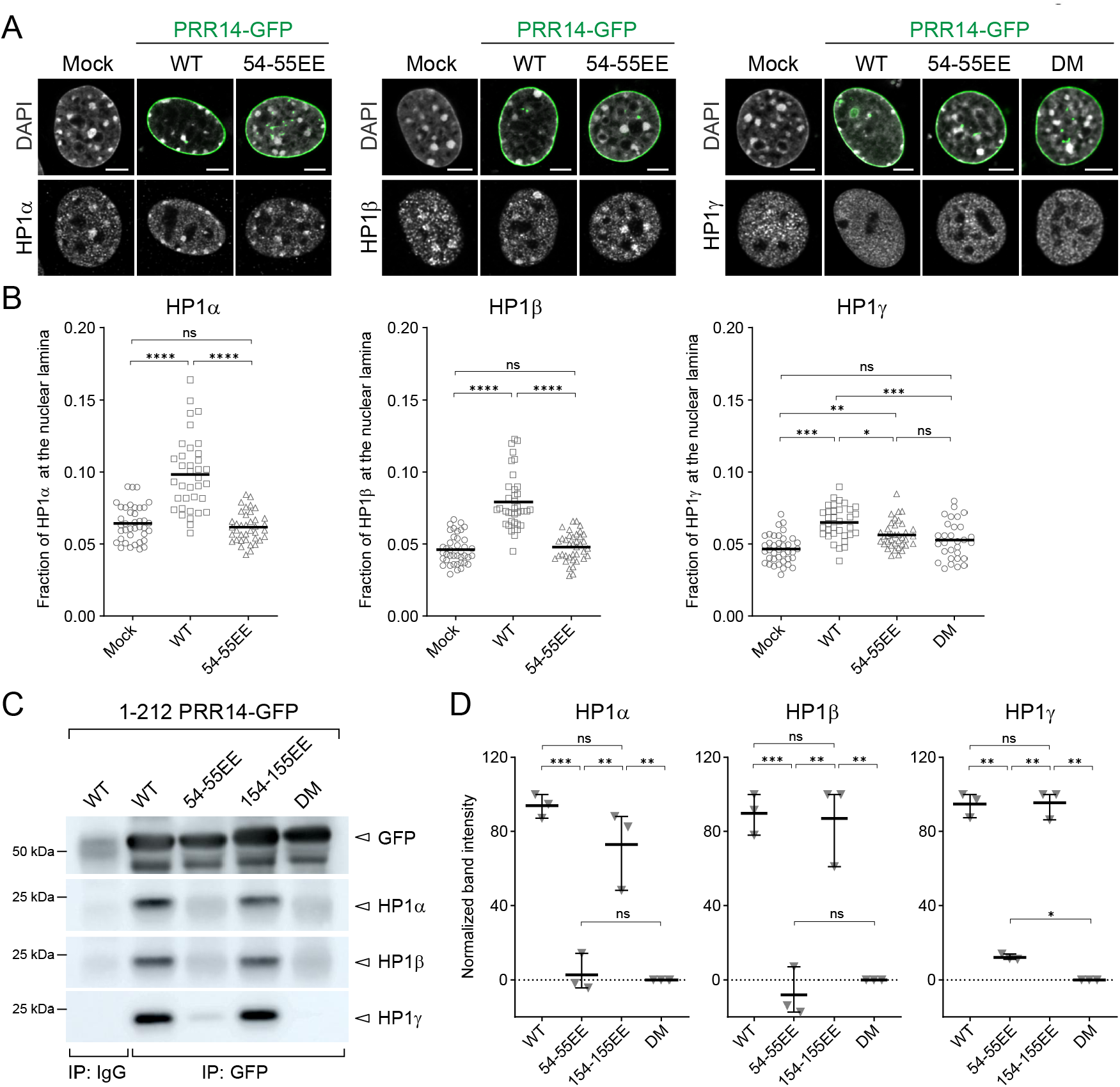
PRR14 tethers chromatin to the nuclear lamina primarily through interactions with HP1α and HP1β. **(A)** Representative confocal images of murine NIH/3T3 cells mock transformed or expressing GFP-PRR14 WT or mutant constructs (green) and stained for HP1 isoforms (red) as indicated. DAPI counterstain shown in gray. **(B)** Dot plots show ratio of indicated HP1 protein signal at the nuclear lamina for murine NIH/3T3 cells expressing GFP-PRR14 WT or mutant constructs as indicated. **(C)** Lysates of 293T cells expressing indicated GFP-PRR14 (1-212) constructs were immunoprecipitated (IP) with anti-GFP antibody and analyzed by Western blotting using antibodies against the indicated HP1 isoform. IgG serves as an isotype control for non-specific antibody interactions. **(D)** Dot plots show normalized intensities of indicated HP1 signal after immunoprecipitation with anti-GFP antibodies. n = 3 independent immunoprecipitations. Lines on dot plots show average values. Whiskers show range. Lines on dot plots show average values. n ≥ 30 cells per condition. Statistical analysis was performed using ANOVA Kruskal-Wallis test with Dunn’s multiple comparisons for panel B and paired ANOVA test with Geisser-Greenhouse correction for panel D; ****p < 0.0001, ***p < 0.001, **p < 0.01, *p < 0.05, ns: not significant. Scale bars 5μm.

We then compared the repositioning of HP1 isoforms by different mutant forms of PRR14. PRR14 54-55EE overexpression did not reposition HP1α or HP1β (Fig. 5A-B), consistent with the minimal interaction of this mutant with H3K9me3-modified heterochromatin presented previously (see Fig. 1). PRR14 54-55EE overexpression had an observable but not statistically significant effect on HP1γ (Fig. 5A-B), and no relocalization of HP1γ was seen in cells overexpressing the DM construct (Fig. 5A-B). Given the measurable effect of WT PRR14 overexpression on HP1γ, we used siRNA knockdown of HP1γ to determine whether HP1γ contributes to PRR14-chromatin interactions. In this assay, we observed a statistically significant decrease of PRR14 association with chromatin (Fig. S7). Overall, our data demonstrate that PRR14 HP1-binding site 1 (LAVVL, aa 52-56) binds all 3 HP1 isoforms and is essential for heterochromatin tethering to the nuclear lamina (Figs. 1 and 5). PRR14 HP1-binding site 2 (LVVML, aa 153-157) may contribute to strengthen overall PRR14 association with chromatin (Fig. 4).

We next assayed the PRR14 interactions with HP1 isoforms through immunoprecipitation followed by Western blot. Given the low solubility of full-length PRR14, we used PRR14 1-212 fragments to assess the PRR14-HP1 isoform interactions. Using anti-GFP antibodies, GFP-PRR14 was immunoprecipitated from cells expressing WT GFP-PRR14 1-212 and the co-immunoprecipitated proteins were subjected to Western blot analysis of individual HP1 isoforms. All three HP1 isoforms were detected following immunoprecipitation which, consistent with the relocalization experiments, indicated that PRR14 can interact with each of these proteins (Fig. 5C, D, Fig. S8).

To determine whether the two HP1 binding sites of PRR14 showed any differences in terms of HP1 isoform interaction, we quantified the amounts of each HP1 protein that co-precipitated with WT PRR14 1-212 compared with the 54-55EE, 154-155EE and DM forms when expressed in cells at comparable levels (Fig. 5C). The interactions between PRR14 HP1-binding site 1 mutant (54-55EE) and all 3 HP1 isoforms were significantly reduced compared with WT, although a small amount of residual binding to HP1γ was retained (Fig. 5C-D). In this assay, we could not detect any significant difference between WT and PRR14 HP1-binding site 2 (154-155EE) mutant in co-precipitating any of the 3 HP1 isoforms (Fig. 5C-D). Analysis of the double mutant (DM) showed complete lack of HP1-PRR14 interaction (Fig. 5C-D). These results are consistent with a primary role for HP1 binding site 1 as a mediator of PRR14 interaction with all 3 HP1 isoforms, and an apparently small contribution of HP1 binding site 2 to the interaction of PRR14 with HP1.

### PRR14 association with the nuclear lamina is independent of heterochromatin binding

PRR14 association with heterochromatin through HP1 is highly dynamic, as described above (Fig. 4). Next, we sought to characterize PRR14 interactions with the nuclear lamina, which have been shown to occur between the PRR14 lamina-binding domain (LBD) and Lamin A/C (*18*). The relative contributions of PRR14-nuclear lamina and PRR14-heterochromatin binding for nuclear lamina localization were investigated using FRAP. We assayed the dynamics of PRR14 at the nuclear lamina for both the full-length protein and the isolated PRR14 LBD (residues 231-351) (Fig. 6A). Full length WT PRR14 protein exhibited a high level of immobile fraction at the nuclear lamina (55% on average), and the recovery half-time of PRR14 at the nuclear lamina averaged 25 seconds (Fig. 6B-C). By way of comparison, the recovery half-time for Lamin A/C at the lamina has been reported to be from 10 to 60 minutes (*36*), suggesting that PRR14 interactions with the lamina are relatively quite dynamic.

**Figure 6.**
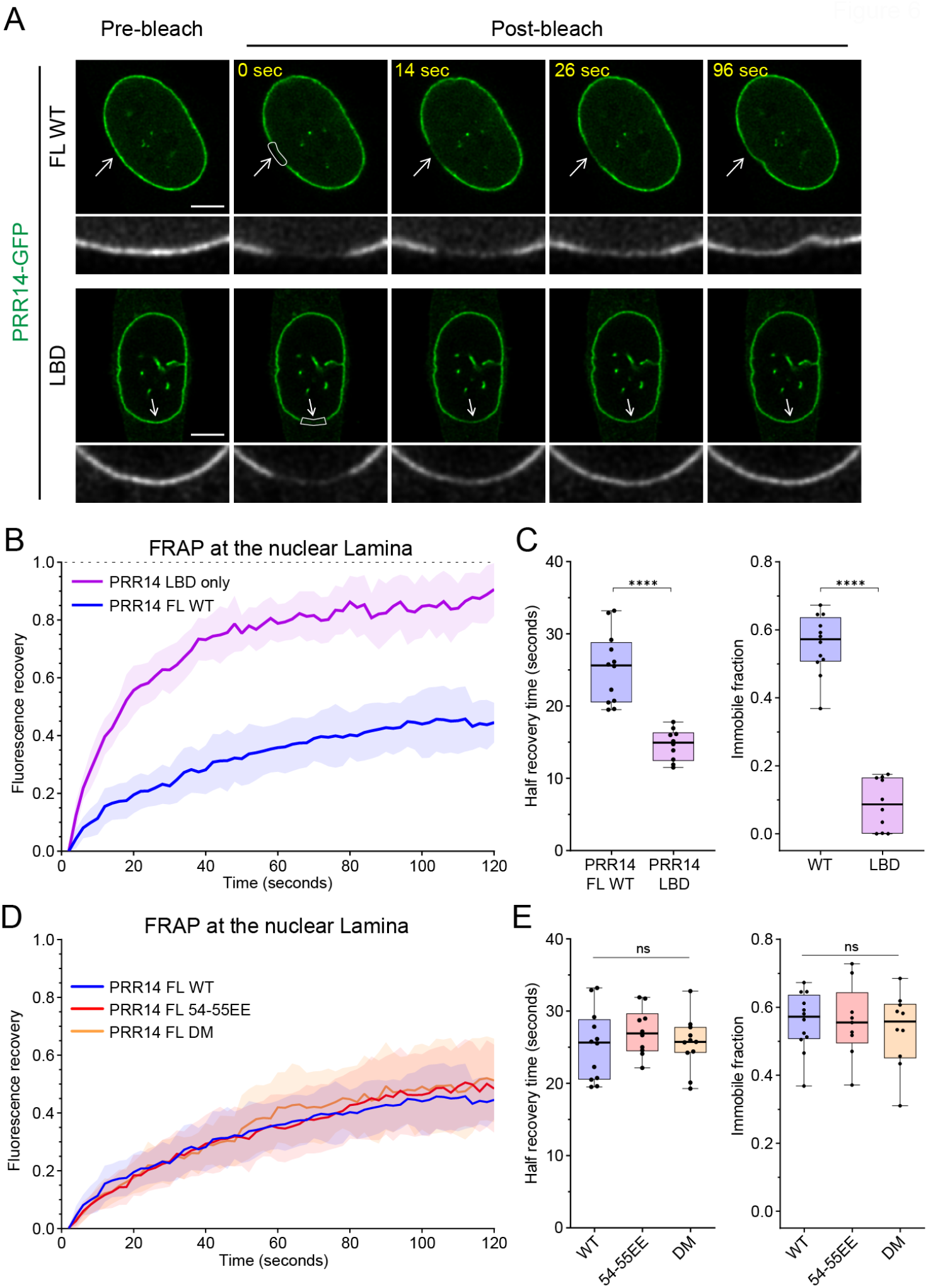
PRR14 association with the nuclear lamina is dynamic and independent of PRR14-heterochromatin binding. **(A)** Representative confocal images of fluorescence recovery after photobleaching (FRAP) assay of WT GFP-PRR14 full-length (PRR14 FL) and lamina-binding domain (LBD) constructs. White boxes indicate bleached area. Grayscale images show magnified bleached areas indicated by white arrows. **(B)** Line graph shows normalized FRAP signal in the areas indicated with boxes in (A) for PRR14 full-length (blue) and PRR14 LBD-only fragment (purple). **(C)** Box plots show distributions of recovery half-time (left) and immobile fraction (right) for indicated constructs. **(D)** Line graph shows normalized FRAP signal in the areas indicated with boxes (Fig. S6) for PRR14 FL WT (blue) and mutant full-length constructs 54-55EE (red), or double mutant (DM; yellow). **(E)** Box plots show distributions of recovery half-time (left) and immobile fraction (right) for indicated constructs. n ≥ 12 cells per condition. Line graphs show mean values with standard deviations displayed as shading. Box plots show median, 25th and 75th percentiles; whiskers show minimum to maximum range. Statistical analysis was performed using Mann-Whitney test and analysis of variance (ANOVA) Kruskal-Wallis test with Dunn’s multiple comparisons. ****p < 0.0001, ns: not significant. Scale bars 5μm.

In contrast to full-length PRR14, the LBD fragment demonstrated measurably different dynamics with only a small immobile fraction (<15%) and an average recovery half-time of 15 seconds (Fig. 6A-C). The LBD fragment lacks both the N-terminal chromatin-binding region and the C-terminal regulatory region. This suggests that the full-length protein is stabilized at the nuclear lamina through additional interactions that result in a significant immobile fraction.

We tested if interaction with HP1 proteins and/or chromatin contribute to PRR14 retention at the nuclear periphery by assaying the dynamics at the nuclear lamina of full-length PRR14 constructs with HP1-binding site mutants (54-55EE and DM), We found no significant differences between WT and these mutant forms of PRR14 in terms of recovery half-time or amount of immobile fraction (Fig. 6D-E, Fig. S9). This suggests that PRR14 dynamics at the nuclear periphery are independent of PRR14-HP1/heterochromatin binding.

Direct comparisons of PRR14 association with heterochromatin (Fig. 4) and with the nuclear lamina (Fig. 6) indicate that PRR14 association with the nuclear periphery is less dynamic than the association of PRR14 with heterochromatin (Fig. S10). Thus, our results support a model in which PRR14 primarily occupies its position at the nuclear lamina where it performs transient attachment to HP1-associated heterochromatin.

### PRR14 at the nuclear lamina creates a surface for heterochromatin anchoring

We used super-resolution microscopy to visualize the location of PRR14 in the nucleus, examining both the nuclear lamina and the nucleoplasm. Localization of PRR14, Lamin A/C and H3K9me3 were examined by Stochastic Optical Reconstruction Microscopy (STORM) and Figure 7 shows representative images of a plane through the middle of the nucleus (Fig. 7A) and a bottom plane at the periphery of the nucleus (Fig. 7B). PRR14 is observed predominantly at the inner nuclear membrane (INM) forming a similar meshwork as Lamin A/C on the INM surface (Fig. 7B), consistent with analyses of our confocal images. A much smaller fraction of PRR14 was observed in the nucleoplasm (Fig. 7A). Nucleoplasmic PRR14 is almost uniformly distributed and we did not observe clusters specific for H3K9me3-modified heterochromatin (Fig. 7A, B). In comparison, H3K9me3 is apparent as aggregated clusters of heterochromatin both at the nuclear lamina and in the nucleoplasm (Fig. 7A, B). No such clustering was observed for PRR14, suggesting that the nucleoplasmic fraction of PRR14 does not strongly associate with heterochromatin away from the nuclear lamina surface. The localization of PRR14 at the nuclear periphery creates a layer of molecules that are capable of binding H3K9me3-modified heterochromatin associated with HP1 as it contacts the nuclear periphery.

**Figure 7.**
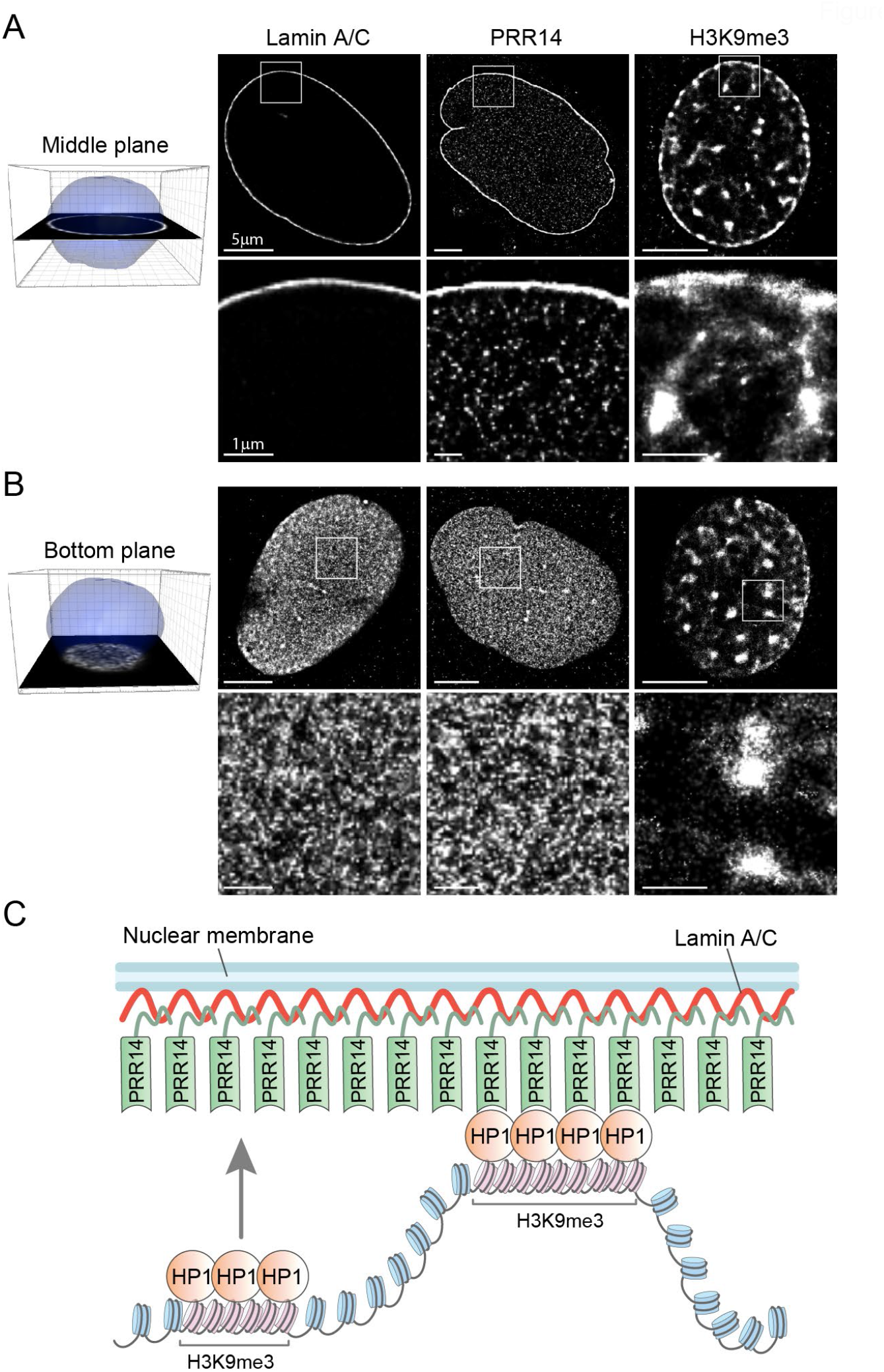
PRR14 is predominantly localized at the nuclear lamina where it creates a surface for anchoring HP1-associated H3K9me3-modified heterochromatin. **(A)** Representative super-resolution stochastic optical reconstruction microscopy (STORM) images showing localization of Lamin A/C, PRR14 and H3K9me3 in the middle plane of nuclei of NIH/3T3 cells. White boxes (top panels) show zoomed areas (bottom panels). **(B)** Representative STORM images showing localization of Lamin A/C, PRR14 and H3K9me3 at the bottom plane of nuclei of NIH/3T3 cells. **(C)** A model illustrating the mechanism for PRR14 tethering H3K9me3-modified chromatin to the nuclear lamina through interaction with HP1. Scale bars: 5μm (top panels) and 1μm (bottom panels).

The results presented here suggest a model in which PRR14 creates a meshwork at the nuclear lamina that organizes H3K9me3-modified heterochromatin at the nuclear periphery through dynamic association with HP1 (Fig. 7C). This PRR14 meshwork might be thought of as a “sticky” surface, or “Velcro”, that allows H3K9me3-modified heterochromatin bound by HP1 to become transiently attached to the nuclear lamina.

## DISCUSSION

In this study, we determined the mechanism of PRR14 in organizing heterochromatin at the nuclear periphery. Results presented here demonstrate that PRR14 overexpression results in repositioning of H3K9me3-modified chromatin to the nuclear lamina. This further supports a function for PRR14 as an organizer of H3K9me3-modified heterochromatin in the nucleus. This appears to be specific to H3K9me3-modified chromatin as we obtained no persuasive evidence that PRR14 alters the localization of H3K9me2-modified chromatin. We have shown that PRR14 acts through its interactions with heterochromatin protein 1 (HP1) to tether chromatin to the nuclear lamina, primarily via HP1α and HP1β isoforms, although our experiments indicate that PPR14 can bind the HP1γ isoform as well. We have also mapped a second HP1-binding motif at the N-terminal end of the PRR14 protein. This site is not sufficient for independent chromatin tethering, but may function to strengthen PRR14 association with chromatin or promote the dense nucleosome compaction characteristic of heterochromatin. Analysis of the dynamics of PRR14 association with the nuclear lamina and chromatin, combined with super-resolution imaging of PRR14 localization in the nucleus and the ability of PRR14 to reorganize heterochromatin localization independent of mitosis, have led us to propose a refined model of heterochromatin tethering at the nuclear periphery. Our results suggest a mechanism through which PRR14 organizes heterochromatin at the nuclear lamina and provide new insights into our understanding of the organization of heterochromatin at the nuclear periphery. Further studies are needed to determine the cellular mechanisms that might alter PRR14 tethering function to regulate heterochromatin association with the nuclear periphery. Notably, the majority (up to 90%) of over-expressed PRR14 localizes at the nuclear periphery, consistent with our previous studies (*29*) and with the lack of dominant-negative effects observed in cells overexpressing mutant PRR14 constructs. This observation also suggests that the cell can tolerate an excess of PRR14 at the nuclear lamina and might adjust PRR14 levels and thereby chromatin organization.

Only a few chromatin tethers have been described to date (reviewed in (*15*)). The list of widely expressed tethers is limited to LBR and PRR14 in mammals (*16, 18*) and CEC-4 in *C. elegans* (*13*). These tethers are expressed in most cell types and have a similar mechanism of action: all of them specifically interact with chromatin bearing H3K9 methyl-modified histones. PRR14 stands out among known tethers as LBR and CEC-4 are integrated nuclear envelope proteins with restricted localization in the nuclear membrane, while PRR14 is a nuclear lamina-associated protein that has dynamic localization at the nuclear periphery.

A notable feature of PRR14 is the duplication of key functional elements. The protein has conserved residues for two NLS signals, two lamina-association motifs, and two HP1-binding sites (*18, 29*). As is frequently seen in evolution, these duplicated regions are not identical. We have yet to determine whether each of these regions provides functional redundancy, modified function, or has already lost their original function through mutational drift. Such protein redundancy might ensure protein function in multiple cell types as well as additional levels of regulation of protein function(s). Our earliest PRR14 study demonstrated that each NLS is independently sufficient to transport PRR14 into the nucleus, but with lesser efficiency singly than when combined (*18*). Therefore, two NLS sequences might serve to efficiently deliver PRR14 in the nucleus. Similar function is observed for the LBD where each lamina-associated domain is able to position the protein to the nuclear lamina independently, but much less efficiently than that observed when both are intact (*29*). Furthermore, LBD-1 region has been shown to be regulated by phosphorylation whereas this has not been observed for LBD-2 (*29*). Such complex organization of the lamina-binding domain allows for regulation of PRR14 association with the nuclear lamina, thus regulating heterochromatin tethering capabilities of PRR14. In this study, we identified a second HP1-binding motif (LVVML, aa 153-157), located near the first HP1-binding site (LAVVL, aa 52-56) in the N-terminus of the protein. Both motifs are a variation of <u>L</u>x<u>V</u>x<u>L</u> sequence which binds the chromo-shadow domain (CSD-CSD) pocket of HP1 (*37*) and are not identical (Fig. 3A). The first HP1-binding motif (LAVVL) of PRR14 was described previously and experiments with substitution of the key amino acids residues in this motif showed a dramatic reduction of PRR14-chromatin association (*18*). Here, we have demonstrated, through a combination of immunoprecipitation and FRAP imaging, that the <u>L</u>A<u>V</u>V<u>L</u> motif of binding site 1 is primarily responsible for PRR14 chromatin binding and tethering (Figs. 1, 3, and 5), while the second <u>L</u>V<u>V</u>M<u>L</u> motif of binding site 2 has a supportive function in PRR14 association with heterochromatin (Figs. 3-4). Machida et al, recently demonstrated that HP1 dimers can bind H3K9me3 histone modifications at two distinct nucleosomes, and thus function to compact heterochromatin by bringing nucleosomes together (*38*). We speculate that an ability of PRR14 to bind two HP1 dimers could also result in chromatin compaction. If this hypothesis is correct, HP1-bound heterochromatin would undergo further compaction when it interacts with PRR14 at the nuclear periphery. Further, the two HP1-binding sites of PRR14 may have an HP1-locking function, similar to that previously described for SENP7 (*35*), in which two HP1 interaction motifs restrict HP1 mobility at heterochromatin, thereby locking HP1 molecules docked on H3K9me3-modified nucleosomes to promote stable HP1 accumulation. Future studies will be aimed at elucidating the role of HP1 in specifying chromatin regions for localization at the nuclear periphery.

While PRR14 was previously reported to bind HP1α (*18, 39*), here we demonstrate that PRR14 can bind all HP1 isoforms. Tethering to the nuclear periphery in the context of PRR14 overexpression was observed largely for HP1α and HP1β isoforms (Fig. 5B). This agrees with the accepted function for HP1α and HP1β in organizing heterochromatin, in contrast to the HP1γ isoform which has been found in both heterochromatin and euchromatin regions (*24, 25*). Previous studies demonstrated that HP1 isoforms can form heterodimers as well as homodimers (*25, 40, 41*). This feature of HP1 isoforms makes it difficult to determine whether a homodimer of a single isoform or a heterodimer is bound by a single PRR14 protein. Specificity of PRR14 for chromatin binding via HP1 could be further complicated by various HP1 dimers and post-translational modifications of HP1 (reviewed in (*22*)). Some HP1 interactions with other binding partners utilize the same PxVxL-binding pocket or position the HP1 dimer deep inside the nucleosome, thus making HP1 unavailable for interaction with PRR14 (*22*). This agrees with the relatively moderate effect of PRR14 overexpression on HP1 relocalization to the nuclear lamina (Fig. 5). Our results provide evidence that PRR14 functions to organize H3K9me3-modified heterochromatin associated with HP1α/β at the nuclear lamina. While the specific genomic regions that are organized at the nuclear periphery by PRR14 are currently unknown, such specificity could be cell type-specific and dictated by a combination of multiple factors including relative levels of HP1 isoforms and PRR14.

Our data demonstrate that PRR14 tethers heterochromatin regions enriched for the H3K9me3 histone modification, which is known to mark silent genes and promoters, but is more abundant in repetitive heterochromatin regions such as major and minor satellites and peripheral heterochromatin, including lamina-associated domains (LADs) (*6, 34, 42*). Localization of heterochromatin at the nuclear periphery is an additional mechanism to efficiently repress expression of cellular genes and repetitive elements. We originally identified PRR14 as an epigenetic repressor and demonstrated that PRR14 knockdown can activate epigenetically silent genes (*30*). We hypothesize that PRR14 can serve as an epigenetic repressor by tethering and thereby silencing genes at the nuclear lamina. To date, the detailed mechanisms of LAD formation and maintenance remain unclear. Both PRR14 and HP1 were recently identified as proteins in the LAD interactome (*43*). PRR14/HP1 likely functions to maintain association of the H3K9me3-marked heterochromatin of LADs with the nuclear lamina. Such PRR14 function could be regulated by PRR14 phosphorylation (*29*), and might be a mechanism for release of genes from LADs.

Recent studies have proposed that histone modifications, specifically H3K9 methylation, can guide chromatin localization in the nucleus (*32, 44*). We have previously suggested a function for PRR14 to reestablish heterochromatin localization at the nuclear lamina at mitotic exit (*31*). We propose that the function of PRR14 in specifically binding H3K9me3 heterochromatin and organizing it at the nuclear periphery is an example of spatial chromatin organization via histone modification and protein tethers.

The results presented here suggest a dynamic model of H3K9me3-modified heterochromatin organization at the nuclear periphery. H3K9me3-modified heterochromatin is observed at perinucleolar heterochromatin and chromocenters, in addition to the nuclear periphery (Figs. 1 and 7). Several studies have described the dynamic behavior of heterochromatin, including that seen at the nuclear periphery, as it is exchanged between the peripheral and perinucleolar compartments (*33*). This dynamic feature of H3K9me3-modified chromatin is consistent with observations that peripheral heterochromatin can be relocated to another heterochromatin compartment after mitosis (*45*). Based on our results, we propose that PRR14 does not form a sustained bond between heterochromatin and the nuclear lamina, but rather creates a surface at the nuclear lamina that is capable of anchoring heterochromatin with which it comes into contact (Fig. 7C).

## MATERIALS AND METHODS

### Cell lines

Mouse NIH/3T3 (ATCC, cat# CRL-1658), human IMR-90 (ATCC, cat# CCL-186) and Lenti-X™ 293T (Takara Bio, cat# 632180) cell lines were cultured in full-DMEM (Corning, cat# 10013CV) supplemented with 10% FBS (Atlanta Biologicals, cat# S11150) and 1x Penicillin-Streptomycin solution (ThermoFisher Scientific/Invitrogen, cat# 15140122). All cultured cell lines were routinely tested for mycoplasma contamination every 3 months using a e-Myco VALID Mycoplasma PCR Detection Kit (iNtRON Biotechnology, Inc., cat# 25239).

### Plasmids

The human wild-type PRR14 expression construct pCMV6-AN-mGFP-PRR14 was previously described (*18, 29*). pCMV6-AN-mGFP-PRR14 wild-type and HP1-binding domain mutants (site 1 mutant (54-55EE), site 2 mutant (154-155EE), and double mutant (54-55EE and 154-155EE)) were made using site-directed mutagenesis. PRR14 1-212 aa fragments (wild-type and the HP1-binding domain mutants, as described above) were amplified from the corresponding pCMV6-AN-mGFP-PRR14 full-length vectors using Q5 Hot-start High-Fidelity Polymerase 2x Mix (New England Biolabs, cat# M0494S) and the following primers containing HindIII and RsrII restriction sites (Forward (HindIII): 5’-GCTTCTCAAGCTTGTACCATCCATGGACTTGCCCGG-3’; Reverse (RsrII): 5’ -TGCGATCGGTCCGCGCTTAGTCTGCAGGCAGAGC-3’). Then, PRR14 1-212 PCR products were sub-cloned into the expression vector pCMV6-AN-mGFP using T7 ligase (New England Biolabs, cat# M0318S) according to the manufacturer’s instructions. Expression levels of PRR14 WT and mutant constructs were observed to be comparable (Fig. S5A).

To make Doxycycline-inducible vectors, PRR14-GFP 1-212 wild-type and the HP1-binding domain site 1 mutant (54-55EE) were amplified from the corresponding pCMV6-AN-mGFP-PRR14 full-length vectors using Q5 Hot-start High-Fidelity Polymerase 2x Mix (New England Biolabs, cat# M0494S) and the following primers containing EcoRI and AgeI restriction sites (Forward (EcoRI): 5’-ACTATAGGGCGGCCGGGAATTCGTCGACT -3’; Reverse (AgeI): 5’ - GATGTCGCGACCGGT TTAGTCTGCAGGCAGAGCAGA -3’). After that, PRR14-GFP 1-212 PCR products were sub-cloned into pLVX-TetOne-Puro (Takara Bio, cat# 631847) vector using T7 ligase (New England Biolabs, cat# M0318S) according to the manufacturer’s protocol.

### Lentivirus production

Lenti-X 293T cells were plated on a 6-well dish and grown to 80% confluent before transfection. To make the lentiviruses containing Doxycycline-inducible PRR14 wt and 54-55EE mutant, 1.2 μg of lentiviral vectors were premixed with 1.2 μg of packaging psPAX2 vector (Addgene, cat# 12260), and 0.75 μg of envelope pMD2.G vector (Addgene, cat# 12259) and transfected into Leni-X 293T cells using Lipofectamine 3000 (ThermoFisher Scientific/Invitrogen, cat# L3000008) transfection reagent. Supernatant containing lentiviruses was harvested 48- and 72-hours post-transfection, passed through 0.45 μm filter to eliminate contamination with Lenti-X 293T cells and cell debris, and pooled together. The lentiviruses were used immediately to infect NIH/3T3 cells.

### Stable cell line production

To generate a stable cell line expressing Doxycycline-inducible WT or 54-55EE mutant of 1-212 PRR14-GFP, NIH/3T3 cells were infected with the corresponding lentiviruses (1/5 – lentivirus/cell culture medium) in the presence of 4 μg/mL polybrene (Sigma-Aldrich, cat# 107689). 4 days post-infection, cells were split and selected for positive clones using 2 μg/mL puromycin for 5 days.

### Transient DNA and siRNA transfection

Lipofectamine 3000 or FuGENE 6 transfection reagents (ThermoFisher Scientific/Invitrogen, cat# L3000008; and Promega, cat# E2691) were used for transient delivery of PRR14 plasmids, in accordance with the manufacturer’s guidelines. Expression levels of GFP-tagged PRR14 full-length WT and mutant constructs, as well as 1-212 PRR14 constructs were observed to be at comparable levels as measured by GFP-signal intensity in flow cytometry (Fig. S11). For super-resolution and confocal imaging cells were plated on 8-well ibidi μ-slides (ibidi, cat# 80826) or on 12×12 mm GOLD SEAL^®^ glass coverslips (Electron Microscopy Sciences, cat# 63786-01) correspondingly, then transfected at 50% confluency and fixed 48 hours post-transfection (for STORM or IF) or imaged live (for FRAP or PRR14/DNA co-localization analysis).

For siRNA transfection, Lipofectamine RNAiMAX transfection reagent (ThermoFisher Scientific/Invitrogen, cat# 13778075) was used according to the manufacturer’s protocol. siGENOME pools of four siRNAs against Mouse Hp1γ or non-targeting siRNA control (Horizon Discovery/Dharmacon, cat# M-044218-01-0005 and D-001206-13-05) at the final concentration of 50 nM were used. Doxycycline-inducible NIH/3T3 cells expressing WT or 54-55 EE mutant of 1-212 PRR14-GFP, described above, were plated on 8-well ibidi μ-slides. The next day, PRR14-GFP expression was induced by adding 2 ug/mL Doxycycline (Sigma-Aldrich, cat# D9891-25g), and at the same time, cells were transfected with the siRNAs. Live cell imaging was performed 72 hours post siRNA transfection and the addition of Doxycycline. Right before the imaging, cells were counterstained with 1 μg/mL of Hoechst 33342 (ThermoFisher Scientific/Invitrogen, cat# 62249) for 10 min. The imaging was performed using a 1x Live Cell Imaging Solution (ThermoFisher Scientific/Invitrogen, cat# A14291DJ) and in a temperature and humidity-controlled microscope chamber set at 37ºC.

### Cell cycle synchronization by a double thymidine block

Cells were synchronized at the G1/S border using a double thymidine block as described previously (*46*). In brief, NIH/3T3 cells were plated on 12×12 mm coverslips in 6-well plates (ThermoFisher Scientific, cat# 140675). The next day cells were pre-treated with 2 mM Thymidine (Sigma-Aldrich, cat# T9250-1G) and 5 hours later transfected with pCMV6-AN-mGFP-PRR14-WT or pCMV6-AN-mGFP-PRR14-site 1 mutant (54-55EE) using FuGENE 6 transfection reagent in the presence of 2 mM Thymidine in the culture medium to arrest cells in G1/S phase. Thirteen hours post-transfection, Thymidine was removed from the medium and the cells were “released” for 9 hours by adding fresh medium to initiate cell-cycle progression. To increase the percentage of G1/S arrested cells, a second round of 2mM Thymidine treatment was performed for another 13 hours, and after that the cells were either kept with Thymidine for additional 12 hours (“Arrested Cells” in Fig. 2) or released and fixed 12 hours post-release (“Cycling Cells” in Fig. 2).

### Flow Cytometry

NIH/3T3 and IMR-90 cells were transfected with PRR14-GFP constructs as described above. At 48 hours post-transfection, live cells were collected and analyzed using BD Accuri C6 flow cytometer (Becton, Dickinson and Company) to assess levels of PRR14-GFP signal intensity.

For the thymidine block experiment, NIH/3T3 cells were trypsinized using 1x Trypsin-EDTA (Invitrogen, cat# 25200056), washed two times in PBS by centrifugation, and fixed with 70% ice-cold ethanol. The cells were stored at -20°C in ethanol for 1 week, then washed twice in PBS by centrifugation, stained with 1 μg/mL final concentration of propidium iodide (Sigma-Aldrich, cat# P4864) for 20 min and processed for flow cytometry/cell cycle analysis.

### Immunofluorescence

Immunofluorescence assays were performed as described previously (*29, 44*). NIH/3T3 and IMR-90 cells were fixed with 2% paraformaldehyde (PFA) (Electron Microscopy Sciences, cat# 15710) for 8-10 min at room temperature, washed 3 times with DPBS (Gibco, cat# 14190-136), and then permeabilized with 0.25% Triton X-100 (ThermoFisher Scientific/Invitrogen, cat# 28314) for 10 min. Then, cells were washed 3 times with DPBS for 5 min and blocked in 1% BSA (Sigma-Aldrich, cat# A4503) in PBS-T (DPBS with 0.05% Tween 20, pH 7.4 (ThermoFisher Scientific/Invitrogen, cat# 28320)) for 60 min. Next, primary antibodies diluted in 1% BSA/TBS-T were added for 1 h, then samples were washed three times with PBS-T for 5 min and incubated with secondary antibodies in 1% BSA/TBS-T for 60 min followed by washing twice with PBS-T and one time with PBS for 5 min. Samples were mounted using Duolink^®^ In Situ Mounting Medium with DAPI (Sigma-Aldrich, cat# DUO82040-5ML). All steps were performed at room temperature.

### Image acquisition and analysis

All confocal immunofluorescent images were taken using a Leica SP8 laser scanning confocal system using 63X/1.40 HC PL APO CS2 objective and HyD detectors in the standard mode with 100% gain. 3D images of the nuclei middle Z-planes were taken as Z-stacks with 0.1 μm intervals with a range of 1 μm per nucleus. Confocal 3D images were deconvoluted using Huygens Professional software by utilizing the microscope parameters, standard PSF and automatic settings for background estimation. Image analysis was performed using ImageJ software (National Institutes of Health, MD). The fraction of the immunofluorescence signal at the nuclear lamina was determined as proportion of signal at the nuclear lamina, measured using a mask of the nuclear lamina signal, to the total signal in the nucleus. Analysis of PRR14 colocalization with heterochromatin was performed using Colocalization Threshold tool in ImageJ with automatic parameters. Pearson’s above threshold coefficient was measured.

### Fluorescence recovery after photobleaching

FRAP imaging was performed on a Leica SP8 laser scanning confocal system equipped with a temperature control incubation chamber and an 63X/1.40 HC PL APO CS2 objective. NIH/3T3 cells were plated on 8-well ibidi μ-slides and transfected with GFP-tagged human PRR14 constructs. FRAP was performed 48 hours post-transfection using LAS X software (Leica Microsystems, Buffalo Grove, IL) with a build-in FRAP wizard at 488 nm laser and a PMT detector. Live cells were imaged using a 1x Live Cell Imaging Solution (ThermoFisher Scientific/Invitrogen, cat# A14291DJ) and in a temperature and humidity-controlled microscope chamber set at 37ºC. FRAP assays for PRR14 1-212aa constructs (Fig. 4) were performed on the same day and under the same conditions. Cells expressing WT and 154-155EE constructs were plated on the same 8-well ibidi μ-slide to minimize sample-to-sample variations. FRAP assays for PRR14 full-length constructs (Fig. 6) were also performed on the same day and under the same conditions. Cells expressing WT, 54-55EE, 154-155EE and LBD constructs were plated on the same 8-well ibidi μ-slide to minimize sample-to-sample variations. For the benefit of data representation, fluorescent recovery curves of WT and LBD, and WT and mutants were presented as two panels (Fig. 6B and 6D), where WT sample is the same.

Briefly, a region of interest (ROI) was bleached for about 4 seconds, then recovery of GFP-PRR14 signal was measured for 40-120 seconds at 0.75-1.5s intervals. Fluorescence intensity of the bleached ROI and total fluorescence intensity of the nucleus was measured using ImageJ software (National Institutes of Health, MD). Out of focus frames were excluded from the analysis. To compensate for the inherent baseline photobleaching over time, the overall recovery signal was normalized to the inherent photobleaching rate of whole nucleus regions. Fluorescent recovery curve fitting and extension was performed in GraphPad Prism 9 software (GraphPad Software, Inc.), as well as calculation of a recovery half-time and an immobile fraction. Immobile fraction was calculated as a complement to 1 of fluorescent recovery plateau/maximum recovery.

### Super-resolution microscopy

Super-resolution imaging was performed using Stochastic Optical Reconstruction Microscopy (STORM). Images were obtained using ONI Nanoimager (ONI Inc., CA). Cells were plated on 8-well ibidi μ-slides and transfected with pCMV6-AN-mGFP-PRR14. Lamin A/C and H3K9me3 staining was performed on untransfected cells. 48 hours post-transfection, samples were fixed, immunostained as described above, and kept in DPBS until image acquisition. Secondary antibodies were conjugated with Alexa 647 dye. Imaging acquisition was performed in fresh imaging buffer (50 mM Tris-HCl, pH 8.0, 10 mM NaCl, 10% (w/v) glucose (Sigma-Aldrich, cat#G8270), 1.5 mg MEA (Sigma-Aldrich, cat#30070), 170 AU Glucose oxidase (Sigma-Aldrich, cat#G2133), 1400 AU Catalase (Sigma-Aldrich, cat#C40)). Images were processed using ONI Nanoimager™ Software version: 1.16 (ONI Inc., CA).

### Immunoprecipitation experiments

GFP-tagged PRR14 1-212 fragments (wild-type, site 1 mutant (54-55EE), site 2 mutant (154-155EE), and DM (54-55EE and 154-155EE)) were transfected into Lenti-X 293T cells and used to assess interactions with HP1 isoforms. Protein A/G Magnetic Beads (ThermoFisher Scientific, cat# 88802) preincubated overnight at +4C° with anti-GFP antibodies or Normal Rabbit IgG were used for pull-down of GFP-tagged proteins or as a mock pull-down. 48 hours post-transfection, cells were washed in PBS, then scraped off the plates in 500 μL ice-cold NP-40 lysis buffer (VWR Life Science, cat# J619-500ML) containing protease and phosphatase inhibitors (Roche, cat# 11697498001 and cat# 4906845001) and 1x benzonase (Sigma-Aldrich, cat# E1014-5KU). The lysates were then incubated for 30 min at +4C° with constant agitation and cleared by centrifugation at 12,000 rpm for 20 min at +4C°. The supernatants were collected and mixed with GFP antibody pre-conjugated beads, which were pre-washed with NP-40 lysis buffer with protease/phosphatase inhibitors (Roche, cat# 11697498001 and cat# 4906845001) and incubated for 2.5 hours at +4C° with gentle agitation. The beads were collected, washed with lysis buffer, resuspended in lysis buffer containing 1x NuPAGE Sample Reducing Agent (ThermoFisher Scientific/Invitrogen, cat# NP0004), 1x Laemmli Sample Buffer (Bio-Rad, cat# 1610747) and heated for 5 min at 95°C to dissociate immunocomplexes from the beads. The beads were then cleared on a magnet, and the supernatants were collected. Proteins were separated on 4-12% SDS–PAGE gels (ThermoFisher Scientific/Invitrogen, cat# NP0335) and Western blotting of a 0.2 μm nitrocellulose membrane (ThermoFisher Scientific/Invitrogen, cat# LC2000) was performed to detect GFP-PRR14 and HP1 isoforms by probing the membrane with the corresponding antibodies. The membranes were developed using ECL Prime Western Blotting Detection Reagents (Amersham/Cytiva cat #RPN2232).

Image analysis was performed using ImageJ Software. In brief, raw RGB Western Blot images were converted into 8-bit, and the colors were inverted. Each band then was selected using an ImageJ Rectangular Tool, and the signal from the selected areas was measured as a mean intensity value. The mean intensity value of the “double-mutant DM” condition was then subtracted from the values measured for the rest of the experimental conditions. Then, the values corresponding to HP1 proteins were normalized to the values of total IP: GFP-PRR14 signal following with a second normalization to the maximum values within each experimental condition.

### Antibodies

The following primary antibodies were used in this study: anti-H3K9me3 (Abcam, cat# ab8898; IF 1:1000), anti-H3K9me2 (Active Motif, cat# 39239, IF 1:1000), anti-Lamin A/C (Santa Cruz, cat# sc-376248; IF 1:500), anti-Lamin B1 (Abcam, cat# ab16048; IF 1:1000), anti-GFP (Abcam, cat# ab290; WB 1:1000, IP 1:200), Normal Rabbit IgG (Cell Signaling, cat# 2729; IP 1:200), anti-HP1α (Abcam, cat# 77256; IF, WB 1:1000), anti-HP1β (Abcam, cat# ab10478; IF, WB 1:1000), anti-HP1γ (Santa Cruz, sc-398562; IF 1:500, WB 1:1000).

The following secondary antibodies from ThermoFisher Scientific/Invitrogen were used: Donkey anti-Rabbit Alexa Fluor 568 (cat# A10042, IF 1:1000), Donkey anti-Mouse Alexa Fluor 488 (cat# A10037, IF 1:1000), Donkey anti-Rabbit Alexa Fluor 647 (cat# A31573, IF 1:1000), Donkey anti-Mouse Alexa Fluor 647 (cat# A31571, IF 1:1000); and HRP-linked anti-Rabbit IgG and HRP-linked anti-Mouse IgG (Cell Signaling, cat# 7074 and 7076; WB 1:1000). IF – immunofluorescence, WB – Western Blot, IP – immunoprecipitation.

### Statistical analysis

For immunofluorescence assays, 10 to 40 individual cells were imaged per condition. These numbers exceed the sample size calculated using power analysis. We considered each imaged individual cell/nucleus as a biological replicate. The numbers of cells per each analysis are indicated in the figure legends. Statistical analysis was performed with GraphPad Prism 9.3.1 software (GraphPad Software, Inc.) using one-way Analysis of Variation (ANOVA) Brown-Forsythe and Welch test with Dunnett’s multiple comparisons (for samples with Gaussian distribution of residuals), paired ANOVA test with Geisser-Greenhouse correction for immunoblot assay, or non-parametric ANOVA Kruskal-Wallis test with Dunn’s multiple comparison test. For comparison of two sample an unpaired non-parametric Student’s t-test (Mann–Whitney test) was used.

## Author Contributions

Conceptualization: A.A.K., R.A.K. and A.P. Methodology: A.A.K., Y.C., C.L.S., and A.P. Formal analysis: A.A.K. and A.P. Investigation: A.A.K., Y.C., and A.P. Visualization: A.A.K. and A.P. Data curation: A.A.K. and A.P. Resources: R.A.K. Supervision: A.P. Writing (original draft): A.P. Writing (review and editing): A.A.K, R.A.K., C.L.S., and A.P.

## Acknowledgments

We thank Jonathan A. Epstein for support (NIH R35 HL140018) and for discussions and comments on the manuscript; Rajan Jain for critical assessment of the manuscript and for helpful suggestions; Kelly Dunlevy, Jason Wasserman, Jade Wilson, Valentina Medvedeva, and Trinity Pellegrin for help with making certain constructs; Andrea Stout from the Penn CDB Microscopy Core for help with confocal imaging; Melike Lakadamyali for help with super-resolution imaging; Center for Engineering and Mechanobiology (CEMB) and NSF Science and Technology Center Pilot Award (CMMI 15-48571) for access to ONI super-resolution microscope; and the Penn Flow Cytometry Core for technical support.

## Data and materials availability

All data needed to evaluate the conclusions in the paper are present in the paper and/or the Supplementary Materials. Raw western blot images are available as source data.

## Supplementary Materials for

**Figure S1.**
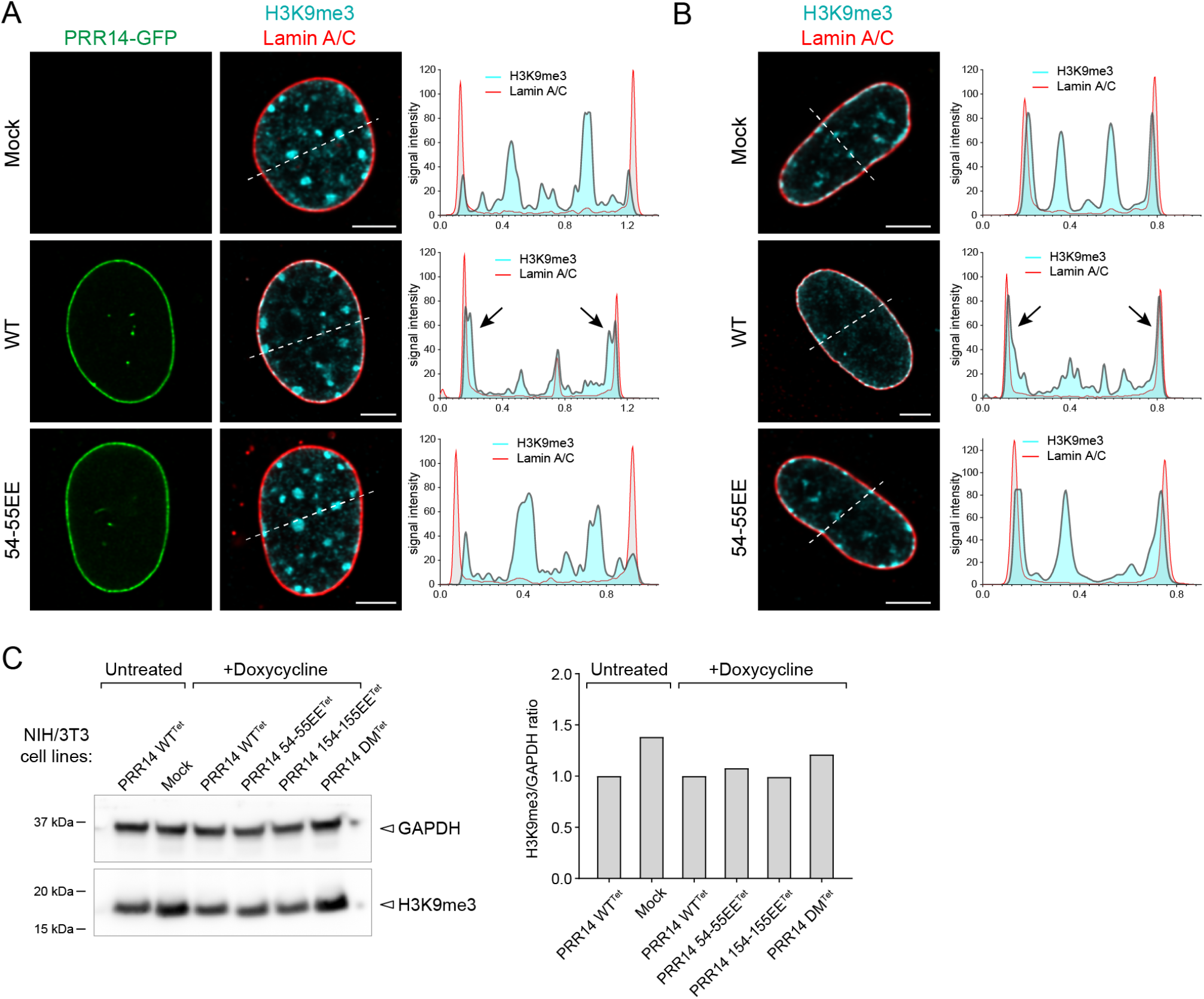
Overexpression of PRR14-GFP resulted in the repositioning of H3K9me3-modified heterochromatin toward the nuclear lamina. Representative confocal images of **(A)** murine NIH/3T3 and **(B)** human IMR-90 cells expressing WT or 54-55EE mutant GFP-tagged PRR14 constructs (green), stained for H3K9me3 (cyan) and Lamin A/C (red) are from Figure 1. Line graphs show signal intensity line profiles of the H3K9me3 (cyan) and Lamin A/C (red) signal across the dotted lines. Arrowheads show repositioning of H3K9me3-modified heterochromatin to the nuclear lamina. Scale bars 5μm. **(C)** Amounts of H3K9me3-modified heterochromatin do not change upon overexpression of PRR14 constructs. Western blot shows H3K9me3 amounts in NIH/3T3 cells (Mock), uninduced and induced NIH/3T3 cells with Tetracycline-inducible vectors coding for the indicated PRR14 constructs (PRR14 WT_Tet_, PRR14 54-55EE_Tet_, PRR14 154-155EE_Tet_, PRR14 DM_Tet_) and GAPDH loading control. Bar graph in right panel shows band intensities as a ratio of H3K9me3 to GAPDH.

**Figure S2.**
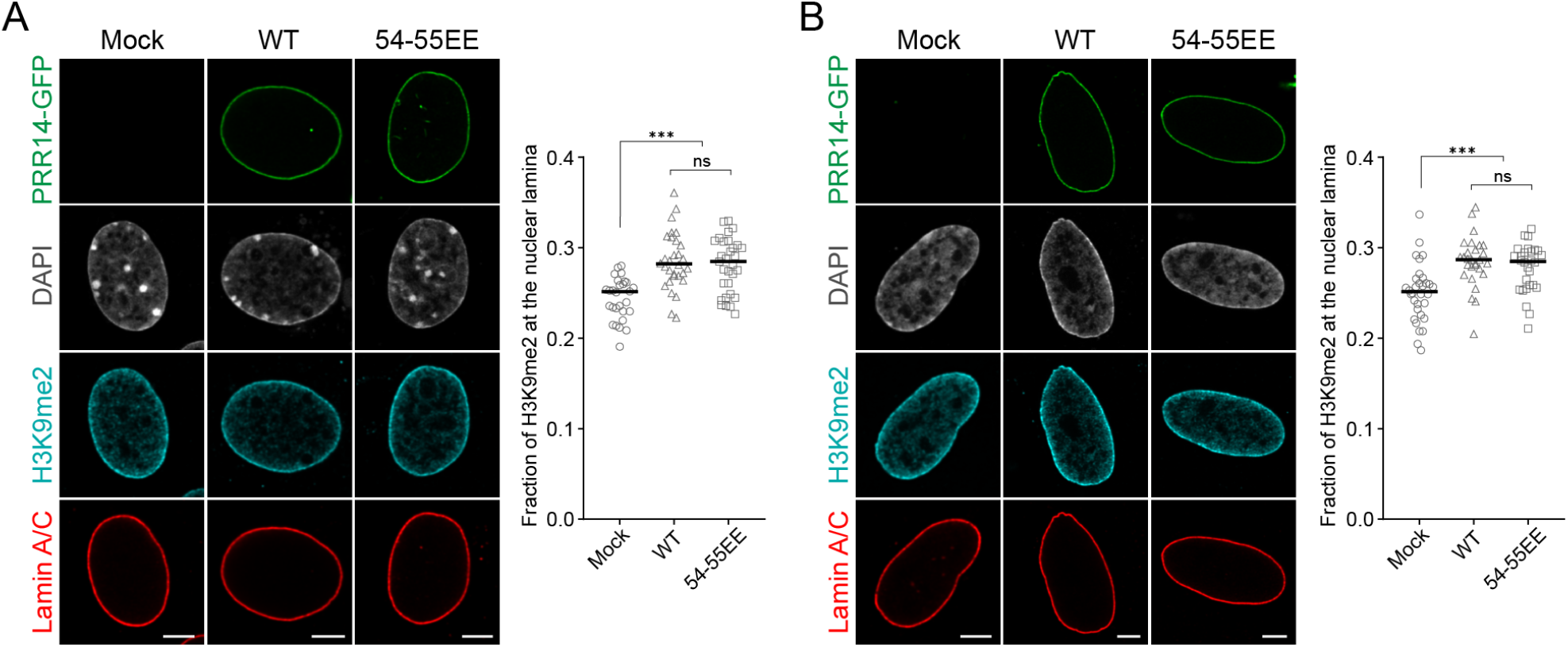
PRR14 overexpression shows no specific tethering effect of H3K9me2-modified heterochromatin localization at the nuclear lamina. While there is a small, but statistically significant effect of PRR14 overexpression on H3K9me2 level at the nuclear lamina, this effect is observed for both WT and the HP1-binding mutant (54-55EE) construct suggesting a chromatin tethering-independent effect. Representative confocal images of **(A)** murine NIH/3T3 and **(B)** human IMR-90 cells expressing WT or 54-55EE mutant GFP-tagged PRR14 constructs (green), stained for H3K9me2 (cyan) and Lamin A/C (red). DAPI counterstain shown in gray. Dot plots show the fraction of H3K9me2 signal at the nuclear lamina. n ≥ 30 cells per condition. Lines on dot plots show median values. Statistical analysis was performed using ANOVA Kruskal-Wallis test with Dunn’s multiple comparisons; ***p < 0.0001, ns: not significant. Scale bars 5μm.

**Figure S3.**
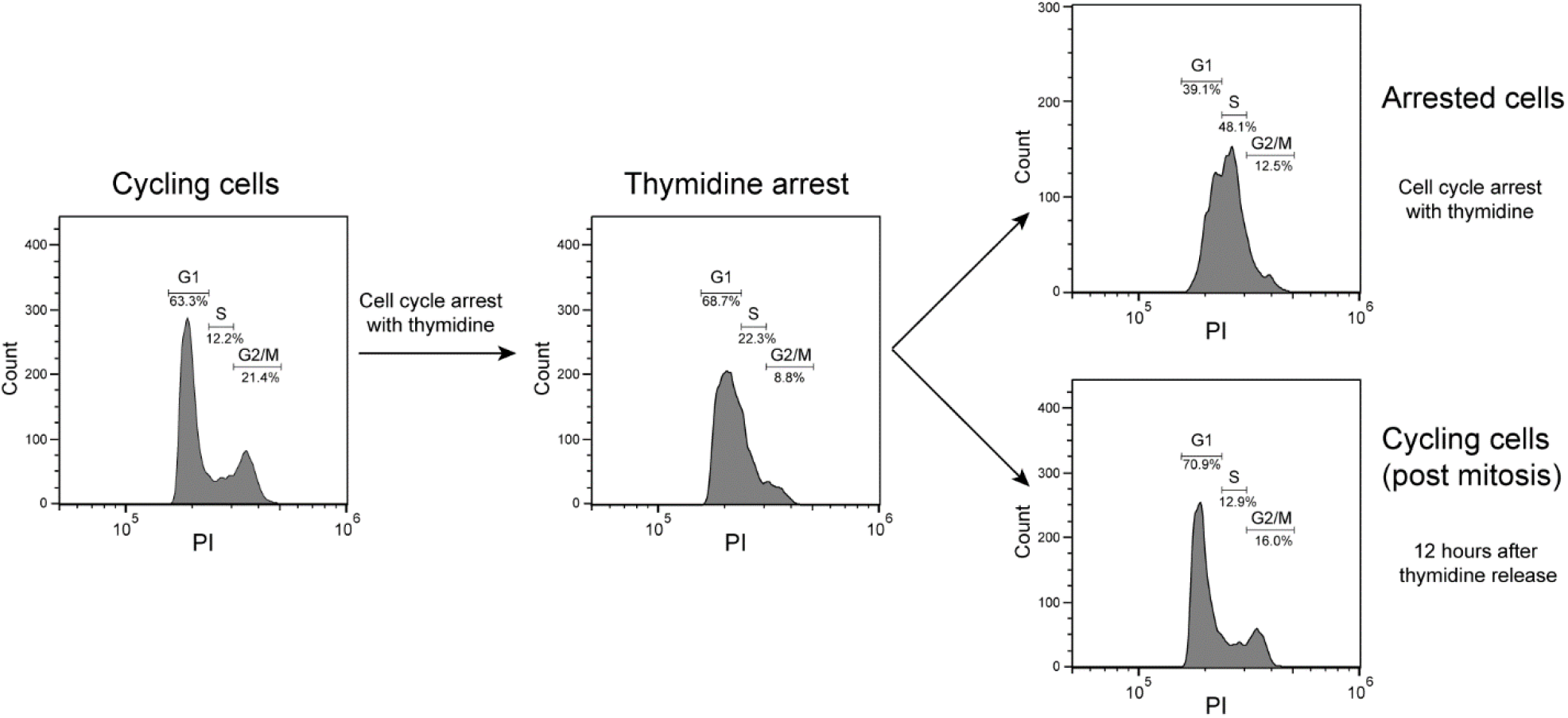
Flow cytometry charts of cell-cycle arrest with thymidine. NIH/3T3 cells transfected with PRR14-GFP were arrested with thymidine to prevent mitosis. Samples were maintained in thymidine to prevent mitosis or released from thymidine block 12 hours before fixation to allow mitosis. Flow cytometry charts show normal cycling cells, thymidine-arrested cells, arrested cells 12 hours post-thymidine treatment, and post-mitotic cells 12 hours after thymidine release. G1, S, and G2/M gates show percentage of cells in indicated cell-cycle stages. PI, propidium iodide.

**Figure S4.**
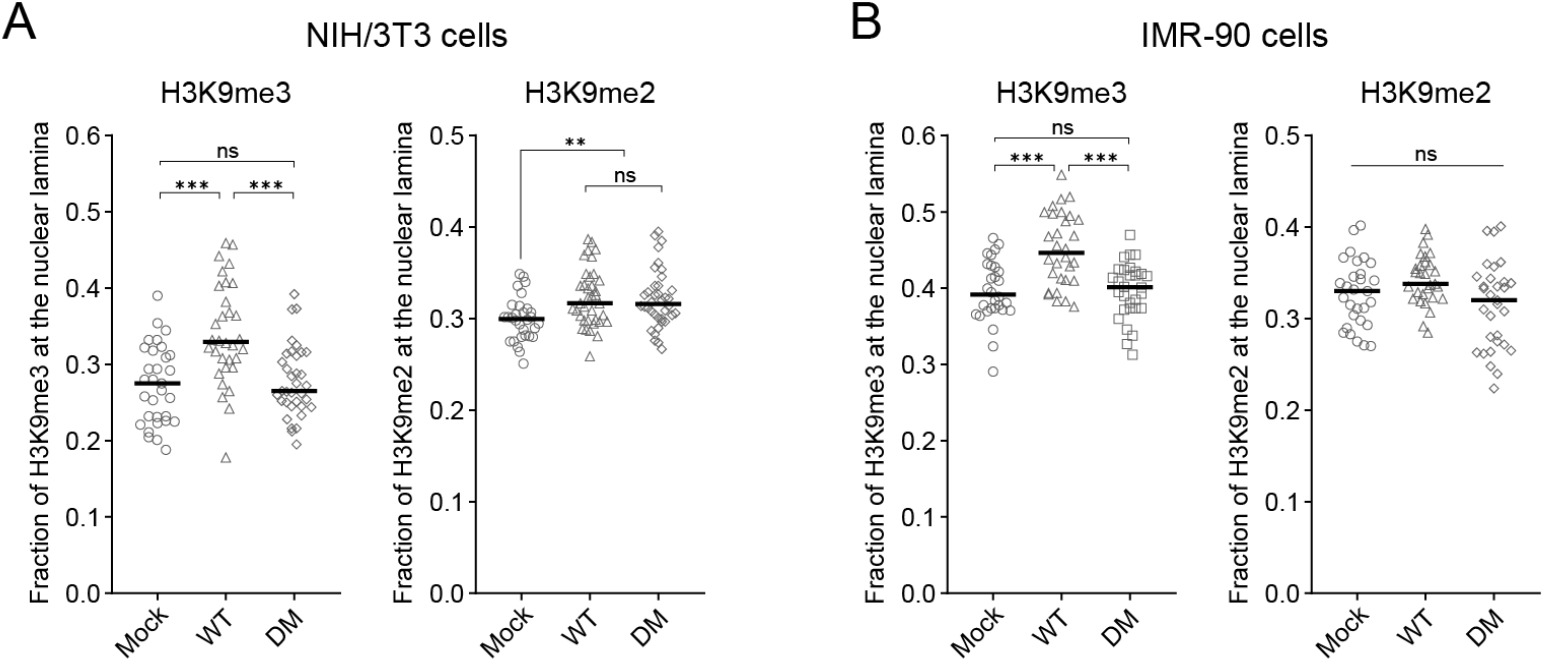
PRR14 specifically organizes H3K9me3-modified but not H3K9me2-modified chromatin at the nuclear periphery. To exclude the possibility that PRR14 HP1-binding site 2 (LVVML) functions to bind and tether H3K9me2 (see figure S1), we performed the same experiment as in Figure 1 and Figure S1 with PRR14 double mutant (DM) construct (54-55EE and 154-155EE). As presented in Figure 1 and Figure S1, we observed a specific tethering of H3K9me3-modified chromatin, but no specific tethering of H3K9me2. This further supports PRR14 specificity for tethering of H3K9me3-modifed chromatin rather than H3K9me2. Observed elevation of H3K9me2-modified chromatin at the nuclear lamina upon PRR14 WT and mutant construct overexpression is not fully reproducible and potentially unrelated to the PRR14 function as a chromatin tether. Dot plots show fractions of H3K9me2- and H3K9me3-marked chromatin at the nuclear lamina in **(A)** NIH/3T3 and (**B**) IMR-90 cells transfected with WT or DM PRR14. n > 35 cells per condition. Statistical analysis was performed using ANOVA Kruskal-Wallis test with Dunn’s multiple comparisons; ***p < 0.001, **p < 0.01, ns: not significant.

**Figure S5.**
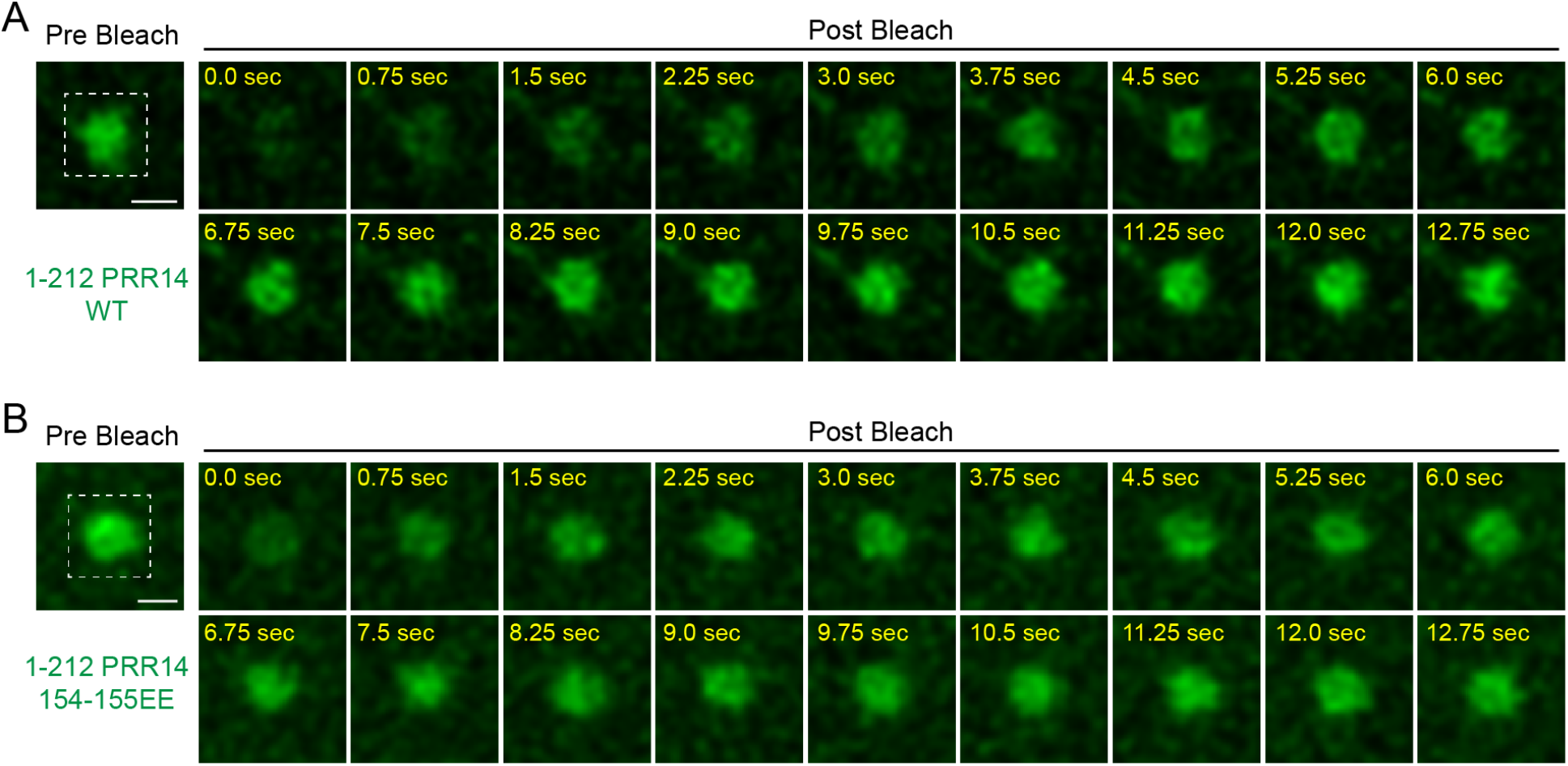
PRR14 association with the nuclear lamina is independent of PRR14-heterochromatin binding. Zoomed regions of representative confocal images shown in Figure 4 of fluorescence recovery after photobleaching (FRAP) assay of WT 1-212 PRR14 and mutant construct 154-155EE. Scale bars 1μm.

**Figure S6.**
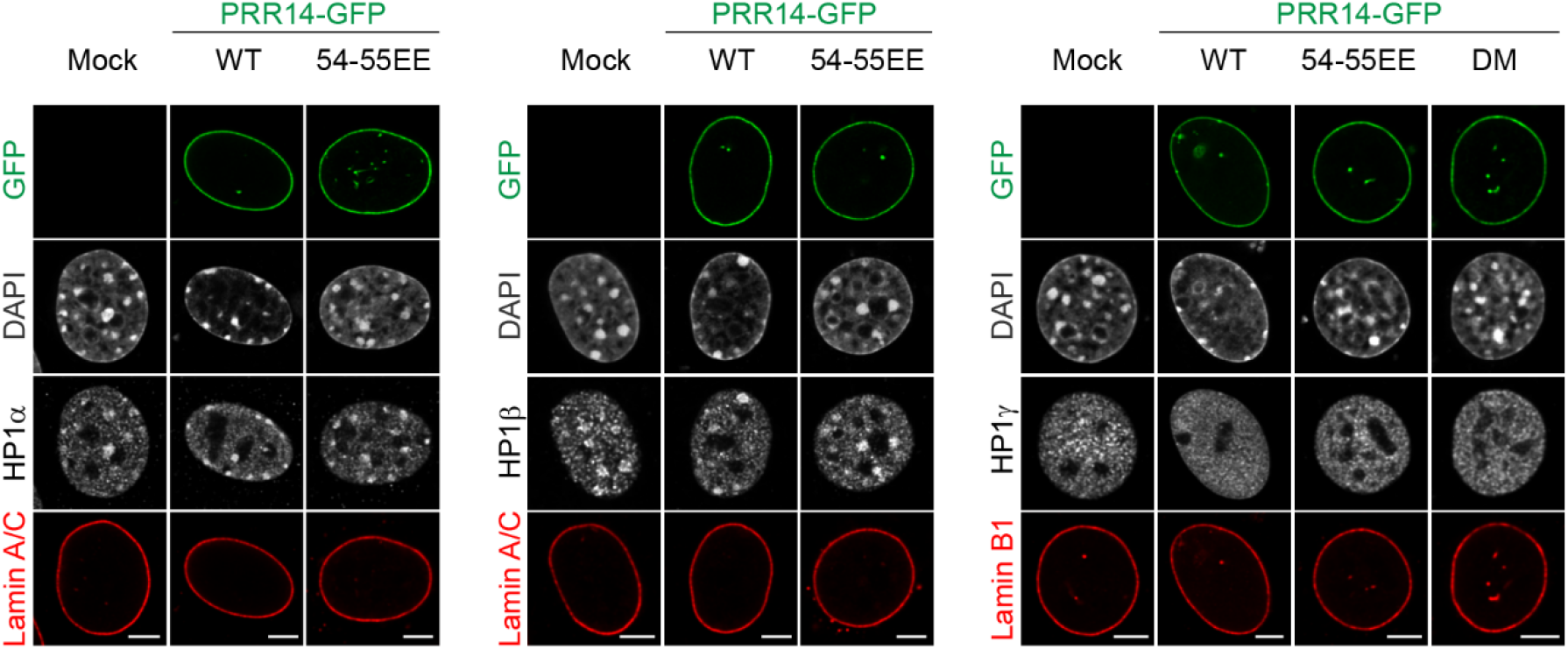
Overexpression of PRR14 results in repositioning of HP1 proteins towards the nuclear periphery. Representative confocal images (from Figure 5) show expression of WT PRR14-GFP or mutant forms and localization of HP1α, HP1β, and HP1γ (in grayscale) in NIH/3T3 nuclei. Scale bars 5μm.

**Figure S7.**
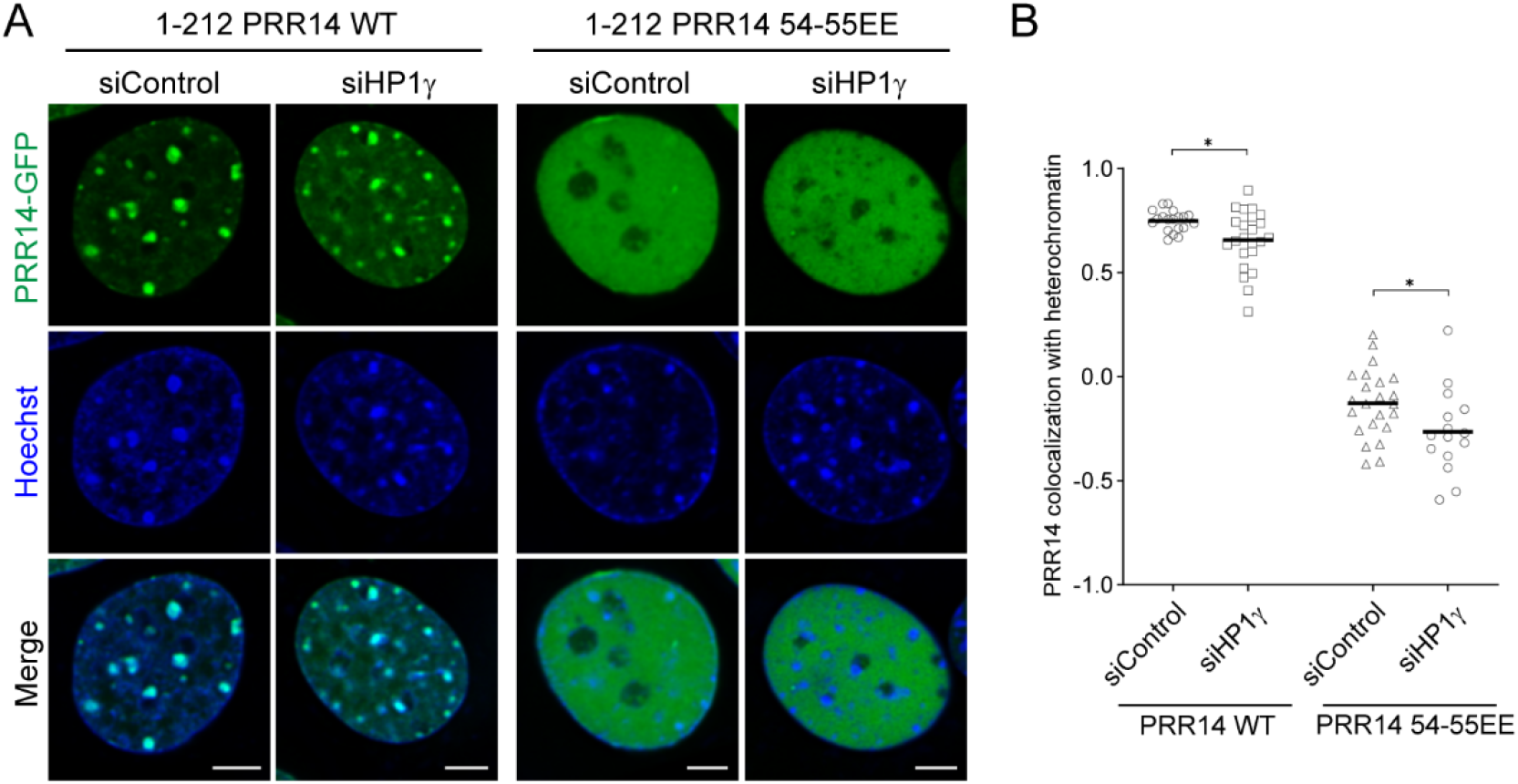
Knockdown of HP1γ decreases PRR14 efficiency for chromatin binding. **(A)** Representative confocal images of murine NIH/3T3 cells expressing indicated WT or mutant GFP-tagged 1-212 PRR14 constructs (green), counterstained with Hoechst (blue). **(B)** Dot plot shows the Pearson’s correlation of Hoechst staining and GFP-PRR14 signal for each PRR14 construct indicating degree of colocalization of PRR14 with heterochromatin regions. n ≥ 15 cells per condition. Lines on the dot plot show median values. Statistical analysis was performed using Mann-Whitney test; *p < 0.05. Scale bars 5μm.

**Figure S8.**
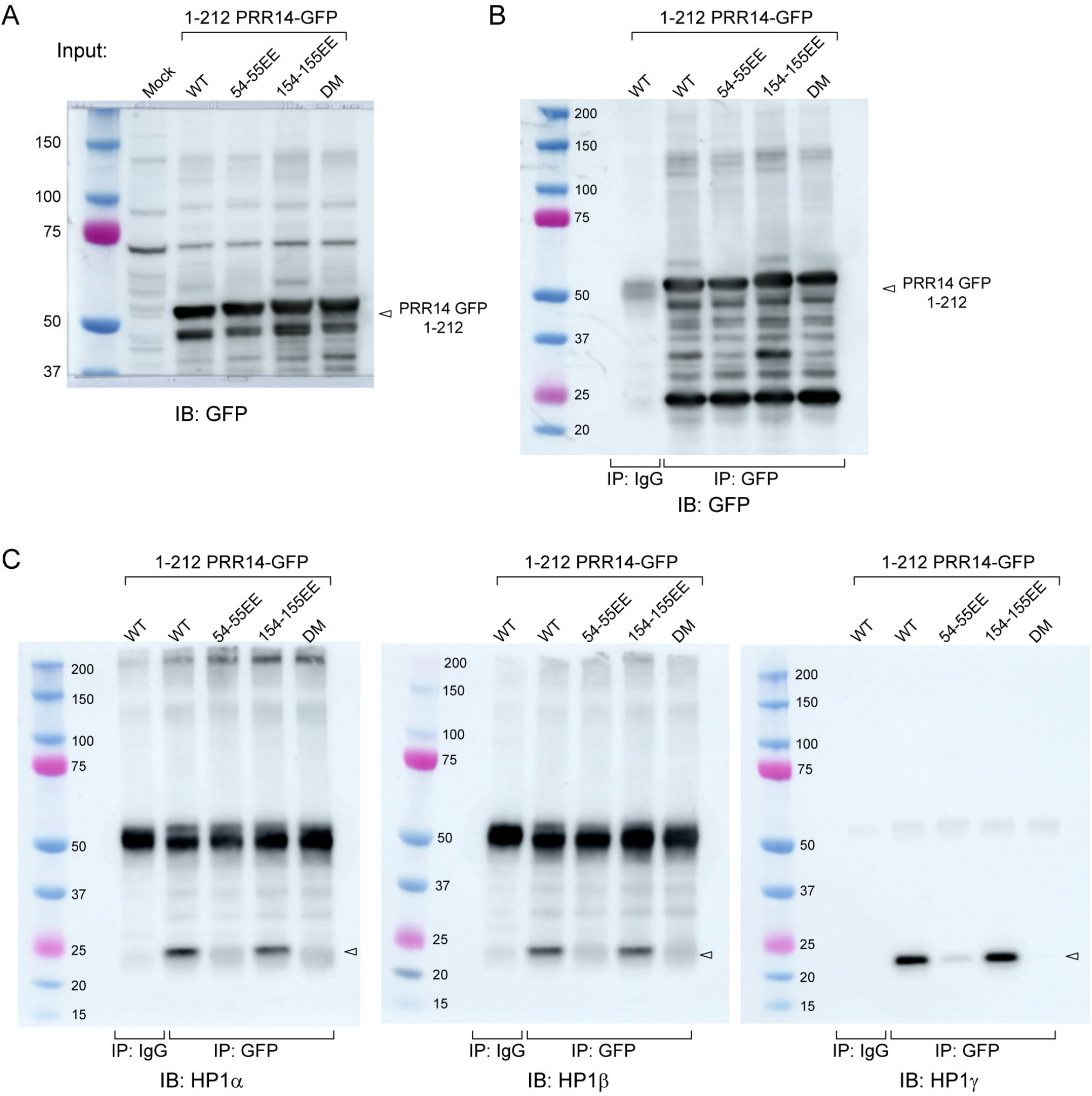
Representative images of Western blot experiments. **(A)** Immuno-blot (IB) shows sample inputs for immunoprecipitation followed by HP1 isoform Western blot experiment with overexpressed GFP-PRR14 1-212 fragment (WT or mutants as indicated). The membrane was immuno-blotted with anti-GFP antibodies. **(B)** Immunoprecipitation (IP) of indicated samples with anti-IgG or anti-GFP antibody, immuno-blotted with anti-GFP antibody. **(C)** Immunoprecipitation (IP) of indicated samples with anti-IgG or anti-GFP antibody, immuno-blotted (IB) with anti-HP1α (left), anti-HP1β (middle) and anti-HP1γ (right) antibodies. Arrowheads show HP1 protein bands.

**Figure S9.**
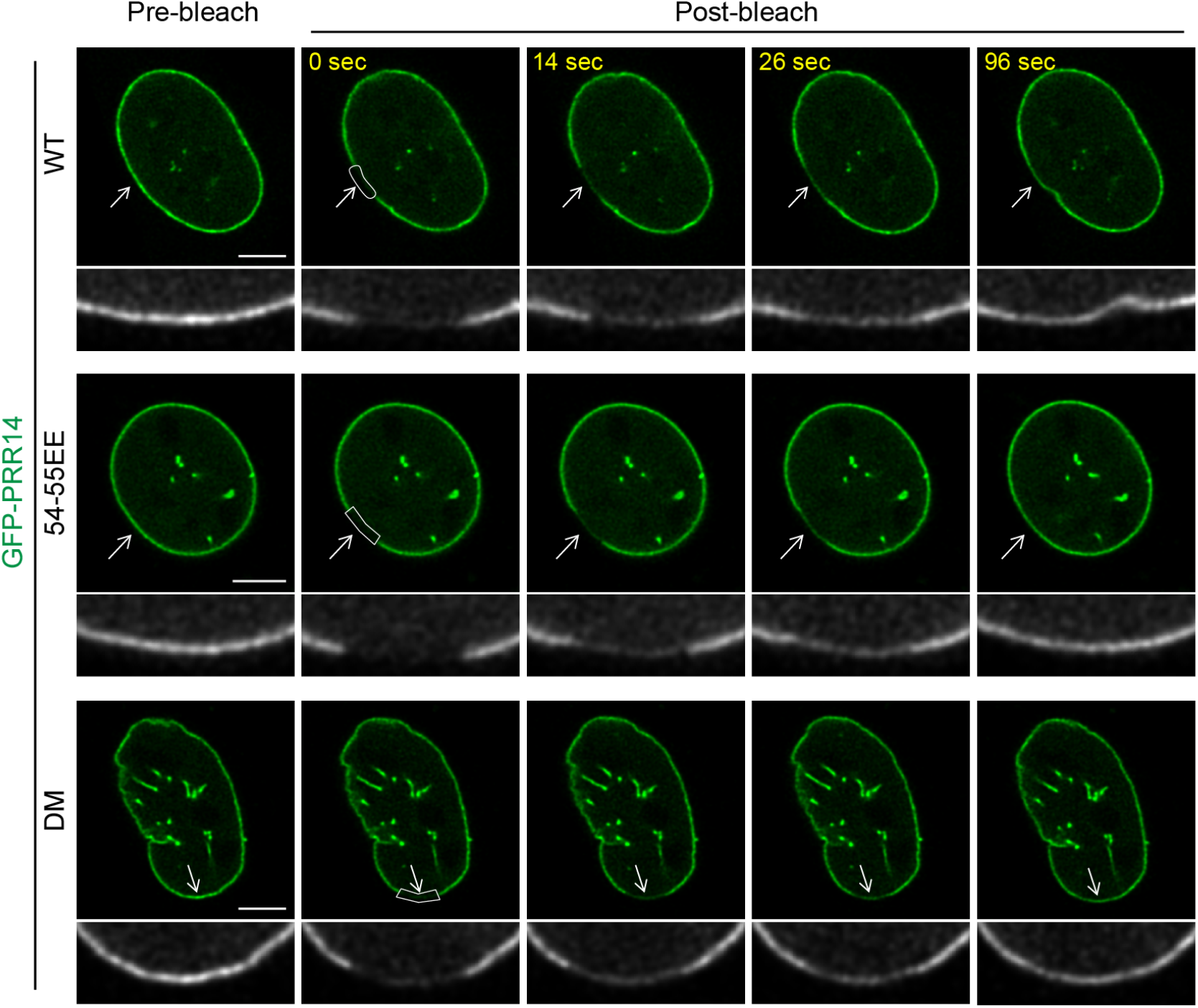
PRR14 association with the nuclear lamina is independent of PRR14-heterochromatin binding. Representative confocal images of fluorescence recovery after photobleaching (FRAP) assay of WT full-length PRR14-GFP and mutant constructs in NIH/3T3 cells. DM, double mutant (54-55EE and 154-155EE). Grayscale images show magnified bleached areas, white arrows. Scale bars 5μm.

**Figure S10.**
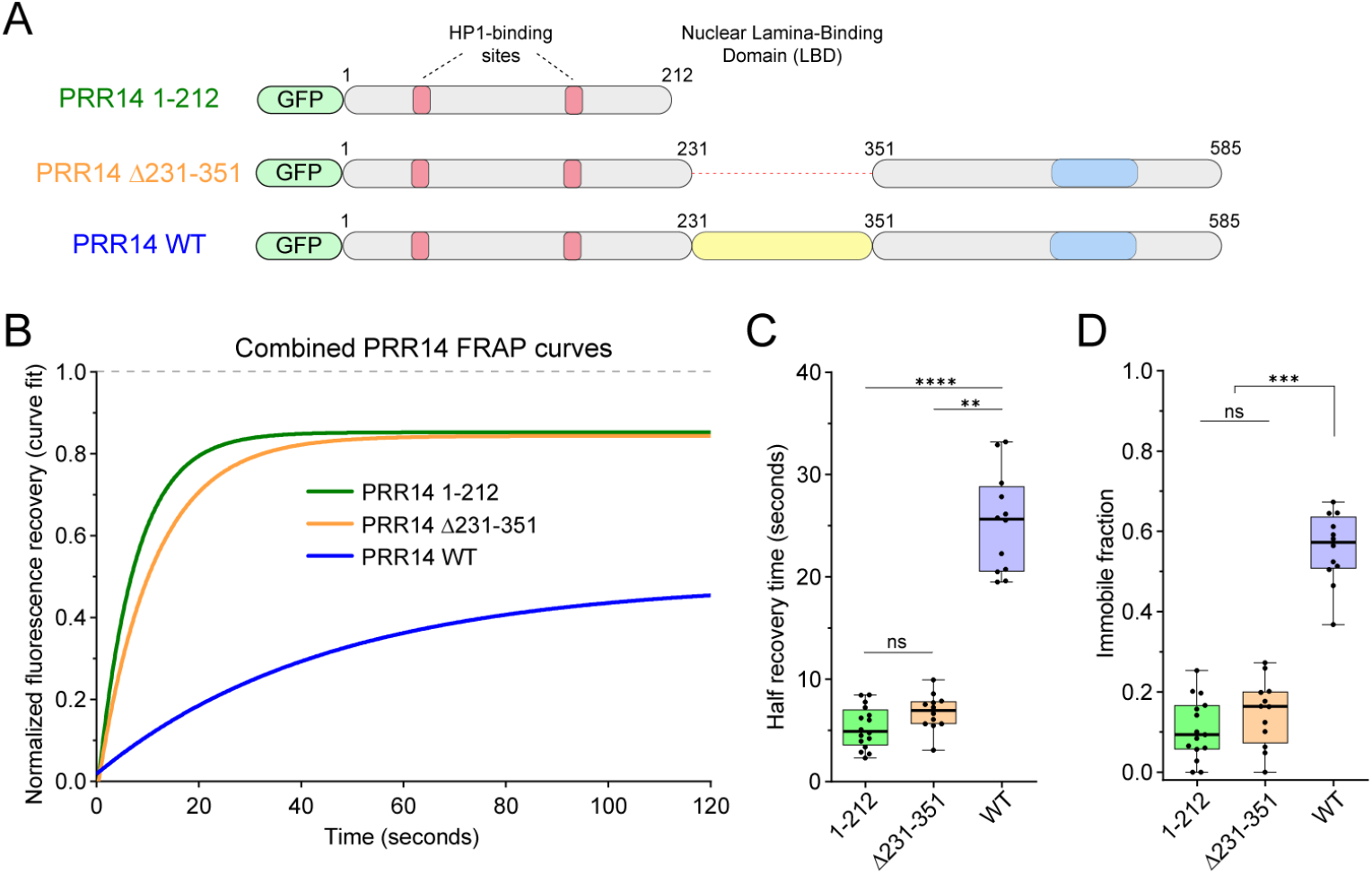
PRR14 shows rapid exchange with heterochromatin, compared to less dynamic full-length WT PRR14 association with the nuclear lamina. To determine whether the rapid PRR14 dynamics of interaction with heterochromatin result from the smaller size of the GFP-PRR14 1-212 fragment (450 aa) used in the FRAP assay compared to the full-length PRR14 (823 aa), we performed a FRAP assay with a GFP-PRR14 construct that carries a deletion of LBD (PRR14 Δ231-351) which is close in size (703 aa) to the full-length protein. We demonstrated that both PRR14 1-212 and Δ213-351 fragments that lack interaction with the nuclear lamina, show similar dynamics with minor differences compared to full-length PRR14 protein. **(A)** Schematic representation of PRR14 fragments including only heterochromatin-binding domain (PRR14 1-212), a fragment with deletion of the lamina-binding domain (PRR14 Δ231-351), and full-length wild-type protein (PRR14 WT). **(B)** Line graph shows extended curve fits of normalized fluorescent recovery over time after photobleaching of indicated PRR14 constructs. **(C)** Box plot shows distributions of recovery half-times for indicated PRR14 constructs. **(D)** Box plot shows distributions of immobile fractions for indicated constructs. n ≥ 12 cells per condition. Box plots show median, 25th and 75th percentiles. Whiskers show minimum to maximum range. Statistical analysis was performed using ANOVA Kruskal-Wallis test with Dunn’s multiple comparisons. ****p < 0.0001, ***p < 0.001, **p < 0.01, ns: not significant.

**Figure S11.**
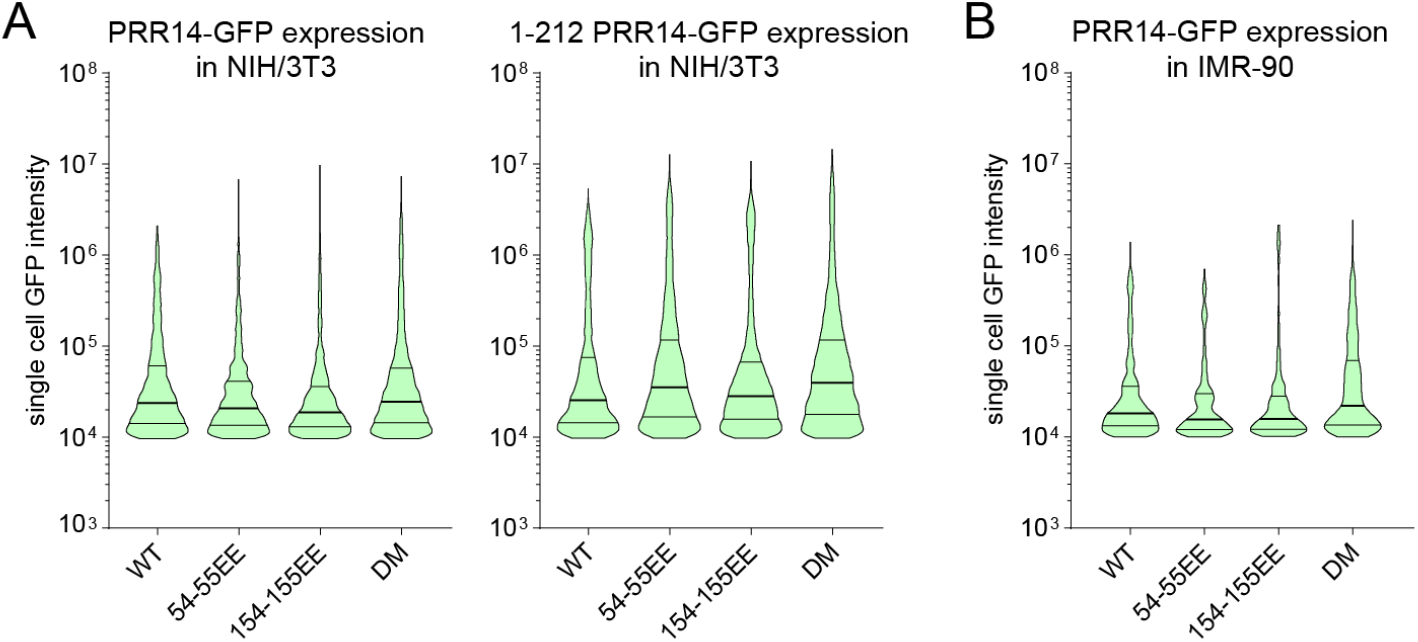
Expression levels of GFP-tagged PRR14 constructs in NIH/3T3 and IMR-90 cells. Minimal expression differences were observed between PRR14 constructs which did not correspond with the observed effects on H3K9me3-modified chromatin localization. Violin plots show GFP intensities of individual cells expressing indicated GFP-tagged full-length and 1-212 PRR14 constructs in **(A)** NIH/3T3 and **(B)** IMR-90 cells. Lines of violin plots show the median and the interquartile range.

